# Assessing biosynthetic gene cluster diversity in a multipartite nutritional symbiosis between herbivorous turtle ants and conserved gut symbionts

**DOI:** 10.1101/2020.05.05.072835

**Authors:** Anaïs Chanson, Corrie S. Moreau, Christophe Duplais

**Affiliations:** Université de Guyane, UMR EcoFoG, AgroParisTech, CNRS, Cirad, INRAE, Université des Antilles, Kourou, France; Departments of Entomology and Ecology & Evolutionary Biology, Cornell University, Ithaca, NY, USA; CNRS, UMR EcoFoG, AgroParisTech, Cirad, INRAE, Université des Antilles, Université de Guyane, Kourou, France

## Abstract

In insect-microbe nutritional symbioses the symbiont supplements the low nutrient diet of the host by producing amino acids and vitamins, and degrading lignin or polysaccharides. In multipartite mutualisms composed of multiple symbionts from different taxonomical orders, it has been suggested that in addition to the genes involved in the nutritional symbiosis the symbionts maintain genes responsible for the production of metabolites putatively playing a role in the maintenance and interaction of the bacterial communities living in close proximity. To test this hypothesis we investigated the diversity of biosynthetic gene clusters (BGCs) in the genomes and metagenomes of obligate gut symbionts associated with the herbivorous turtle ants (genus: *Cephalotes*). We studied 17 *Cephalotes* species collected across several geographical areas to reveal that (i) mining bacterial metagenomes and genomes provides complementary results demonstrating the robustness of this approach with metagenomic data, (ii) symbiotic gut bacteria have a high diversity of BGCs which is correlated with host geography but not host phylogeny, (iii) the majority of the BGCs comes from the bacteria involved in the nutritional symbiosis supporting conserved metabolic functions for colonization, communication and competition in the gut environment, (iv) phylogenetic analysis of arylpolyene, polyketide (PK), and siderophore shows high similarity between BGCs of a single symbiont across different ant host species, while non-ribosomal peptide (NRP) shows high similarity between BGCs from different bacterial orders within a single host species suggesting multiple mechanisms for genome evolution of these obligate mutualistic gut bacteria.

## Introduction

Bacterial symbiosis is widespread among insects and has shaped the evolution of their hosts^1,2^. The symbiotic interactions between bacterial communities inside the host, and between bacteria and hosts, relies massively on metabolites, metabolic pathways and on the enzymes that regulate metabolic flux. Among mutualistic bacteria the nutritional symbionts supplement a large diversity of nutrients to the diet of their insect hosts which often rely on nutritionally limited food sources. In the case of strictly blood-sucking ticks, intracellular bacterial symbionts provide vitamin B to the host^3,4^. Xylophagous termites rely on a diverse community of gut bacteria to degrade the lignocellulose from wood and also fix nitrogen for their hosts.^5^ Herbivorous aphids and ants feeding on nitrogen-poor diets depend on gut bacteria to recycle nitrogen from food waste and contribute to the biosynthesis of essential and non-essential amino acids^6,7^. Bees have a conserved gut bacteria community able to degrade polysaccharides^8^. Herbivorous beetles have bacteriocytes hosting bacterial symbionts which enrich the host metabolism with aromatic amino acid to support cuticle formation^9^. Herbivorous turtle ants also need their gut bacterial symbionts to support normal cuticle formation (Duplais et al. in review).

Another type of microbial symbiosis involves defensive mechanisms where bacterial symbionts enhance insect resistance to a variety of natural enemies including microbes, fungi or nematodes^10,11^. Bacteria can produce a wide range of secondary metabolites which are not required for the immediate survival of the bacteria, but serve important functions in microbial-microbial interactions, such as defense, competition and communication^12,13^. For example, in several insect groups, including beetles, wasps, bees and ants, bacterial symbionts produce a cocktail of antimicrobial compounds to protect the nest^14–16^, eggs^17^ or a mutualistic fungal strain^15,18^.

Bacterial metabolites are produced via specific groups of genes, called biosynthetic gene clusters (BGCs), which are located in close proximity to each other in the bacterial genome. Together, the genes composing these clusters encode for specific enzymatic steps in the biosynthetic pathway of metabolites^19^. Bioinformatics tools have been recently developed to mine bacterial genomes^19,20^ and retrieve the BGCs of different chemical families including arylpolyene, lanthipeptide, non-ribosomal peptide, polyketide and terpene. Metabolites play several roles in bacterial communities for colonizing tissue, communication through quorum sensing, competition using antimicrobial molecules, and nutrient acquisition with siderophore. BGC studies often focus on the genomes of culturable strains from environmental bacteria^21,22^ or host-associated bacteria^23,24^ and can contribute to the discovery of bioactive compounds for medicine. Mining of bacterial metagenomes has also been investigated from environmental samples^25^, however this approach is only starting to be applied to bacterial communities associated within hosts^26^.

The study of BGCs from symbiotic bacteria may help us predict the role of metabolites encoded in symbiont genomes and provide a framework to understand which functional genes are maintained to support bacterial colonization and maintenance within a host. In addition the comparison between bacterial BGCs across different hosts^27^ from different geographic areas^28^ allows us to test what environmental, ecological or phylogenetic factors shape BGC diversity in host-associated bacterial communities.

To investigate the potential drivers of BGCs diversity in insect symbionts, we investigated the association between gut bacteria and herbivorous *Cephalotes* ants. In this nutritional symbiosis the nitrogen-poor diet of ants is supplemented by a core microbiome which recycle urea food waste into amino acids beneficial to the ant^29^. Genomic approaches have revealed the redundancy in functions related to the nitrogen flux in the genome of five conserved bacterial families from the gut of 17 species of *Cephalotes*^7^. Unlike most intra- or extracellular single nutritional symbionts, the *Cephalotes* multipartite gut bacteria mutualism may retain metabolic functions selected not solely for direct benefit to hosts, but for sustaining diverse bacterial community members. To assess if the gut bacteria associated with *Cephalotes* possess BGCs, and if the environment and evolutionary history of the *Cephalotes* are correlated with BGCs diversity, we focused on bacterial genomes and metagenomes from the gut of 17 *Cephalotes* species collected from North and South America^7^. We retrieved a high number of BGCs from both genomes and metagenomes of bacteria associated with *Cephalotes* ants and tested the correlation with geography and host phylogeny. To study the potential role of metabolites in this multipartite mutualism we recorded the bacterial origin of each BGCs to understand their occurrence in gut bacteria involved in this nutritional symbiosis with *Cephalotes.* An analysis of core genes from BGCs (arylpolyene, non-ribosomal peptide (NRP), polyketide (PK) and siderophore) was performed to identify architectural similarity patterns across different symbionts and host species. Our work shows the relevance of genome and metagenome mining in an insect-microbe symbiosis to study the evolution of BGCs in symbiont genomes across the host phylogeny within an ecology and evolutionary framework.

## Material and Methods

### Genomes and metagenomes analysis

The 14 genomes of cultured gut bacteria and 18 metagenomes of *Cephalotes* gut bacteria were obtained from JGI-IMG version 5.0^30^ (Table S1 and S2 respectively) from the previous projects Gs0085494 (“*Cephalotes varians* microbial communities from the Florida Keys, USA”), Gs0114286 (“Symbiotic bacteria isolated from *Cephalotes varians*”), Gs0117930 (“*Cephalotes* ants gut microbiomes”) and Gs0118097 (“Symbiotic bacteria isolated from *Cephalotes rohweri*”)^7^. The metagenomic data used in this study have two metagenomes from the same *C. varians* species. However, for the metagenome of *C. varians* PL010W the sequencing quality is very different from all the other metagenomes (number of reads, GC content), therefore it was excluded in our analysis (Table S2). The metagenomes were analyzed via the software Anvi’o version 5.5^31^ to sort the different bacterial families composing each metagenome into distinct bins. In this analysis, the fasta sequence of a metagenome is used to create a contig database and a profile database. Then, this contig database is visualized and bins are manually created to maximize completeness while minimizing redundancy. Finally, the software CheckM version 1.1^32^ was used to identify the taxonomy of each bin (Figure S1).

### Bacterial biosynthetic gene clusters analysis

The bacterial biosynthetic gene clusters (BGCs) of each genome and each metagenomic bin were analyzed with antiSMASH 5.0^33^ (Figure S1) with the following analysis options: strict detection, and activation of search for KnownClusterBlast, ClusterBlast, SubClusterBlast and Active site finder. BGCs smaller than 5kb were then filtered out of the data and were not included. The taxonomic classification of each cluster was verified to the genus level using the software Blast+^34^. NaPDos^35^ was used to classify the ketosynthase (KS) domain and condensation (C) domain sequences required in the biosynthesis of PK and NRP respectively and to infer the KS and C phylogenies.

### Distance matrix calculations

The published *Cephalotes* phylogeny was retrieved^36^ and the packages ape^37^ and phytools^38^ of the R software version 3.6.1^39^ were used to exclude from this phylogeny the *Cephalotes* species from which no genome or metagenome was available. The cophenetic distance option in the R package stats^39^ was used on this pruned phylogeny to create the *Cephalotes* phylogenetic distance matrix. The cophenetic distance replaces original pairwise distances between the *Cephalotes* species by the computed distances between their clusters. To calculate the metagenomic BGCs distance matrix, a BGC matrix containing the numbers and types of each BGC found in each metagenome was created. Then the distance function (Euclidean method) of the R package stats was used on the BGC matrix to calculate the metagenomic BGCs distance matrix. The Euclidean method was chosen because it calculates the absolute distance between two samples with continuous numerical variables, without removing redundancies.

### Genetic networking construction

The genetic networking of the genomic and metagenomic bacterial BGCs was generated using BiG-SCAPE version 20191011^40^ (Figure S1). The network was constructed with the three following options: “--include_singletons”, “--mix”, and “--cutoffs 1.0”. The resulting similarity matrix was filtered with different thresholds between 0.6 and 0.8^41–43^, and the threshold 0.65 was chosen because it filtered out enough to form different groups while maximizing the number of bacterial BGCs in each network^41,43^. The filtered similarity matrix was visualized with Cytoscape version 3.7.2^44^, using the MCL clusterization algorithm from clusterMaker2^45^ version 1.3.1 with an index value I = 2.0. To create a genetic networking of the bacterial BGCs we the method of Lin et al.^46^ and implemented in the software BiG-SCAPE by Navarro-Munoz et al.^40,47^

### BGCs genomic core gene analysis

Although BGCs are made of many domains, some have been deemed the “genomic core”. The genomic core is a group of one or a few domains within a single BGC, which contain all the required modules needed for the BGC to be functional and are highly conserved across bacteria. The genomic core of four types of BGCs (arylpolyene, NRP, siderophore and T1PK) were analyzed using the CORASON software version 1.0^40^ (Figure S1). First, a database for each type of BGCs was created from all the BGCs of the same type recovered in the genomes and metagenomes, which were obtained from our previous AntiSMASH analysis. Then a BGC from each of our database was chosen as the reference BGC for its database, and a gene from the biosynthetic core of this reference BGC was chosen as the query protein. The choice of the reference BGC and query protein for a database were made taking different variables into account: (1) the reference BGC must be one of the longest BGCs in the database, (2) the query protein must come from a biosynthetic core gene or an additional biosynthetic gene close to the core, and (3) the query protein must be present in at least half of the BGCs in the database. Once a reference BGC and a protein query were selected for a database, the CORASON software was used to determine the genomic core of this type of BGCs, and to infer the phylogenies. If less than half of the BGCs present in a database appear in the phylogeny, or if many BGCs appear more than once in the phylogeny (which may happen if the selected query protein belongs to a biosynthetic gene that can be found multiple times in the same BGC), the reference BGC and query protein previously selected were deemed to be inadequate for the analysis. In this case, the process to choose a reference BGC and query protein were done once more, following the same procedure as before. The reference BGC and query protein selected for the four types of BGCs studied are presented in Table S3.

### Statistical analysis

To test for correlations between the phylogeny and geography of the *Cephalotes* ants and the BGCs found in the bacterial metagenomes the Chi-squared function of the R package stats^48^ was implemented. The *Cephalotes* phylogenetic distance matrix and the bacterial BGCs distance matrix from the metagenome analysis were compared using the Mantel test of the R package ape^37^. The PCoA between the number and types of bacterial BGCs from the metagenome analysis and *Cephalotes* geography were performed using the *prcomp* function of the R package stats and the R package ggplot^49^. The correlations between the number of each type of bacterial BGCs from the metagenome analysis and *Cephalotes* geography were calculated using the one-way ANOVA function with the Tukey’s pairwise of the Past software version 3.25^50^.

## Results

### Mining bacterial genomes

The whole bacterial genomes (N=14) studied herein originated from cultured gut bacteria associated with *Cephalotes varians* and *Cephalotes rohweri*. Among these 14 bacterial genomes, only three genomes did not possess any BGC, while in the others a total of 31 biosynthetic gene clusters (BGCs) were found and the type and length of the 31 BGCs are listed in Table S4. In bacteria associated with *C. varians*, seven types of BGCs were found for a total of 10 BGCs: arylpolyene, beta-lactone, ectoine, nonribosomal peptide (NRP), resorcinol, polyketide type I (T1PK) and terpene. In bacteria associated with *C. rohweri*, six types of BGCs were found for a total of 21 BGCs: arylpolyene, beta-lactone, ladderane, NRP, siderophore and T1PK. In *C. varians* the majority of BGCs originate from Burkholderiales with only one BGC coming from Pseudomonadales symbionts whereas in *C. rohweri* they originate from different bacterial orders (Burkholderiales, Opitutales, Pseudomonadales and Xanthomonadales). The number of bacterial BGCs from *C. varians* and *C. rohweri* were compared by an ANOVA test (Figure S2). Results show that only in the case of NRPs the number of BGCs are statically different between the two ants host (p=0.0036).

### Assessing the quality of metagenomic binning

The analysis of the 17 *Cephalotes* metagenomes resulted in 168 bins and using AntiSMASH a total of 233 BGCs were found in 102 bins out of 168 (Table S5). Completeness of the BGCs ranges from 2.47% to 100%. More than half the bins (87 out of 168) have a predicted completeness level higher than 85%, 66 bins have a completeness level between 50 and 85%, and only 15 bins have a completeness level lower than 50%. The completeness is strongly correlated with the redundancy and number of contigs and genes in each bin (Figure S3A; N=168; PCA analysis, 1^st^ dimension=92.19%, 2^nd^ dimension=7.74%). The length of each bacterial BGC is highly linked to the number of genes in the corresponding BGC (Figure S3B; N=233; Spearman correlation, S=354810, p<2.2e-16, rho=0.832), and moderately linked to the completeness of the bin in which the BGC is found (Figure S3C; N=233; Spearman correlation, S=1579300, p=9.252e-13, rho=0.251). Among the 168 bins, 31 different bacterial orders were identified, and among them 21 bacterial orders possess at least one bacterial genus predicted to have a BGC (Figure S4A). The bacterial families having the highest number of BGCs are Burkholderiales (41 BGCs), Pseudomonadales (41 BGCs), Rhizobiales (41 BGCs), and Xanthomonadales (24 BGCs). The bacterial specificity of each bin has been assessed by analyzing the number of different bacterial orders present in each bin (Figure S4B). A bin is considered specific if it is associated with only one bacterial order. More than half of the bins are specific, and 75% of the bins contain less than two bacterial orders.

### Mining bacterial metagenomes across the *Cephalotes* phylogeny

From the 233 BGCs retrieved in the genomes and metagenomes we identified 20 different types of BGCs: acyl amino acids, arylpolyene, bacteriocin, beta-lactone, butyrolactone, cyclodipeptide (CDP), ectoine, furan, hserlactone, ladderane, lanthipeptide, linear azol(in)e-containing peptides (LAP), non-ribosomal peptide (NRP), phenazine, resorcinol, siderophore, polyketide type I (T1PK), terpene, and thiopeptide. The arylpolyene and NRP BGCs were the more numerous with 66 and 56 respectively retrieved in the metagenomes. The type and length of all the BGCs are presented in Table S6. Some bacterial BGCs found in the metagenomic analysis are also hybrids made of different types of BGCs: arylpolyene-beta-lactone-resorcinol, arylpolyene-ladderane, arylpolyene-resorcinol, and NRP-T1PK (Table S6). Although we cannot rule misassembling during the genome construction, we are confident in the hybrid NRP-T1PK because the sequence of the KS domain correspond to hybrid KS sequence in NaPDoS database. In Figure 1, the most abundant types of BGCs across the different *Cephalotes* species are investigated, representing a total of 202 BGCs originating from 19 different bacterial orders.

**Figure 1.**
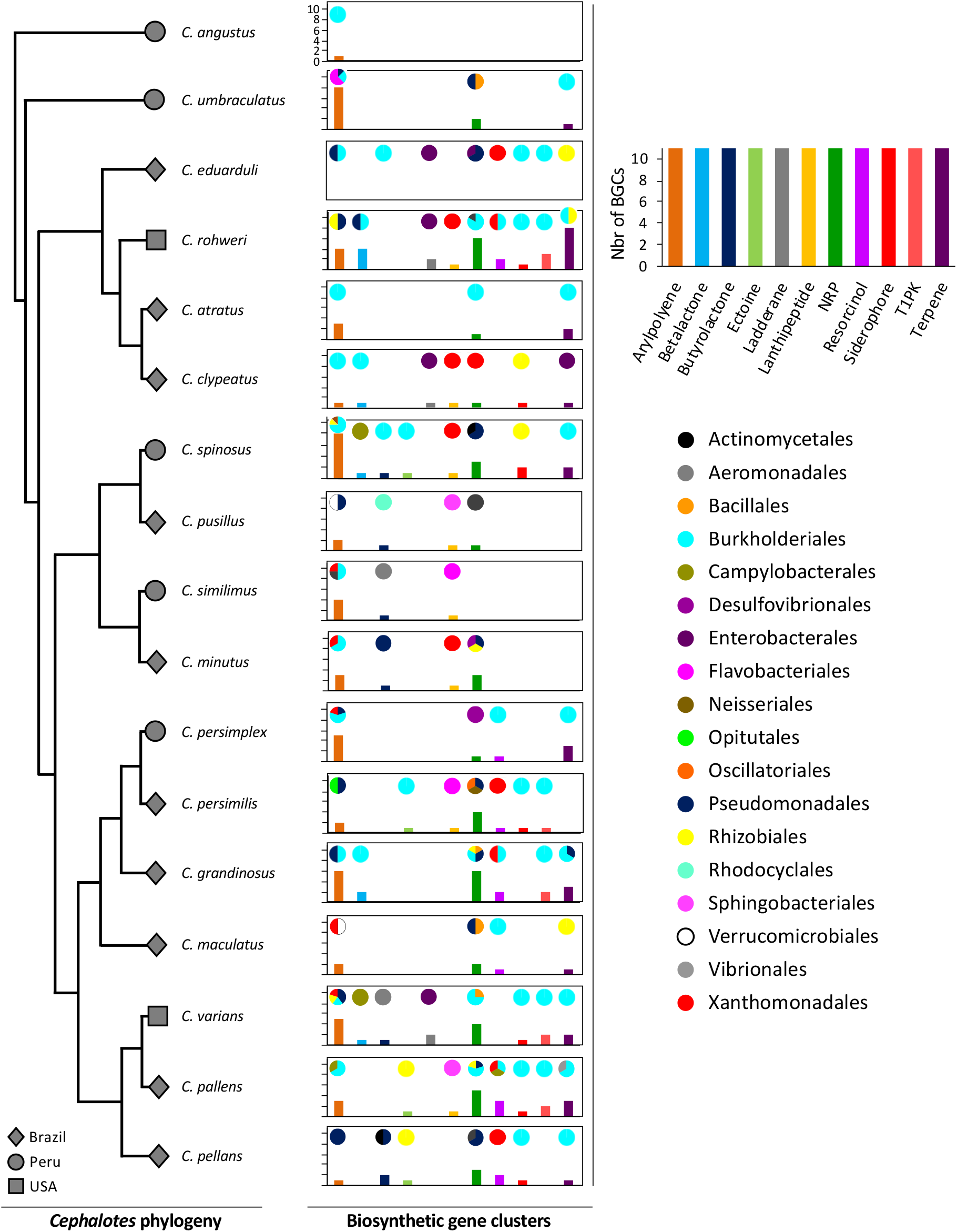
Diversity of biosynthetic gene clusters of bacteria associated with *Cephalotes* turtle ants. The grey symbols represent the country from which an ant originated. The graphs represent the bacterial BGCs found in the metagenomes of each *Cephalotes* species in this study. The bars of the graphs represent the number of BGC of each type found in the metagenomes. The pies represent the proportion of bacterial orders from which each BGC originated.

The 17 species of *Cephalotes* studied here were collected in three different countries, 10 species from Brazil, five species from Peru and two species from the USA. Across these species, a high number of BGCs are found in each species with a mean of ∼14 (Figure 1; Table S6). The lowest number of BGCs was found in *C. angustus* (N=1), the highest number of BGCs was recorded for *C. rohweri* (N=31) and the median number of BGCs across species is 14. The correlations between the diversity of bacterial BGCs and the phylogeny or geography of the *Cephalotes* hosts were assessed (Figure 2). After a Chi-squared test confirmed that the phylogeny and geography of the ants are not correlated (p = 0.558), a Mantel test indicates no correlation between the *Cephalotes* phylogeny and the type and number of bacterial BGCs (Figure 2A; N=17; p=0.205; Mantel test). On the other hand the type and number of bacterial BGCs are statistically different in ants collected in Brazil, Peru and USA (Figure 2B; N=17; p=0.004; ANOVA). A principal coordinates analysis shows that the bacterial BGCs associated with *Cephalotes* coming from the same country group together (Figure 2B; PCoA, 1^st^ axis=51.89%, 2^nd^ axis=25.42%).

**Figure 2.**
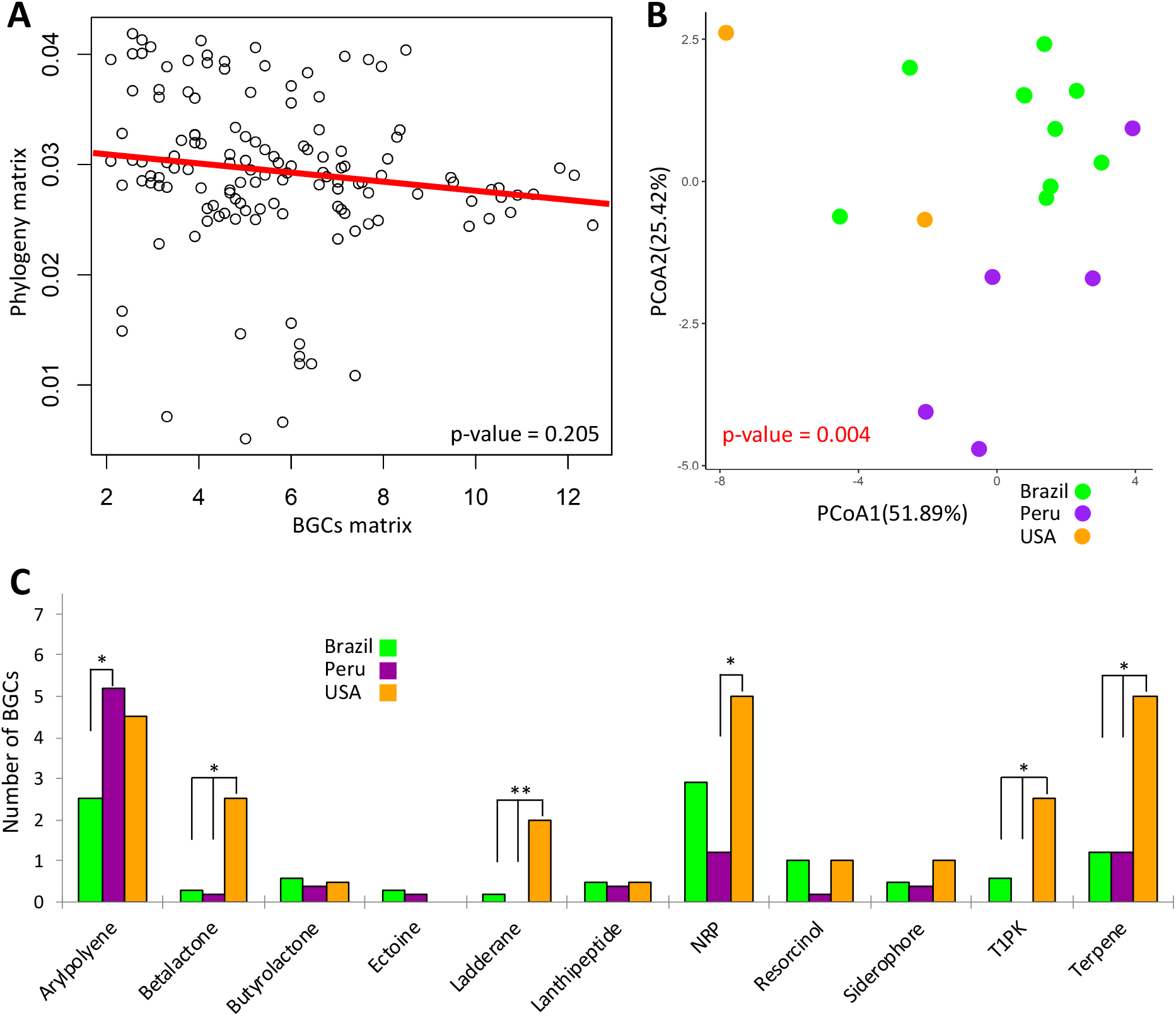
(A) Correlation between the *Cephalotes* phylogeny distance matrix and the bacterial BGCs distance matrix. (B) PCoA of the bacterial BGCs across *Cephalotes* geography. (C) One-way ANOVA test showing the statistical differences between the most abundant bacterial BGCs across *Cephalotes* geography. The symbol * represents p-value < 0.05.

The number of each type of bacterial BGCs associated with *Cephalotes* from different countries were compared using ANOVA (Figure 2C; N=17). Arylpolyene BGCs are more numerous in bacteria of ants from Peru than from Brazil, but not in other comparisons (Peru-Brazil: p=0.035; USA-Peru: p=0.604; USA-Brazil: p=0.409). Beta-lactone, ladderane, T1PK, and terpene BGCs are more numerous in bacteria of ants from the USA compare to ants from Brazil (p=0.047, p=0.014, p=0.049, p=0.048 respectively) and Peru (p=0.037, p=0.010, p=0.038, p=0.040 respectively), while there is no difference between ants from Brazil and Peru for these BGCs (p=0.315, p=0.417, p=0.536, p=1.000 respectively). NRP BGCs are more numerous in bacteria of ants from the USA than in bacteria of ants from Peru (Peru-Brazil: p=0.163; USA-Peru: p=0.046; USA-Brazil: p=0.241). No statistical differences were present in the number of butyrolactone, ectoine, lanthipeptide, resorcinol, and siderophore BGCs in bacteria of ants between the different geographic areas (p=0.720, p=0.560, p=0.760, p=0.880, p=0.620 respectively).

### Genetic networking and BGCs genomic core genes phylogenies

To assess the genetic architectural diversity of BGCs (number and type of gene, gene order, gene domain, and BGC length) we created a genetic network of the BGCs found in the genomes and metagenomes of *Cephalotes*-associated bacteria (Figure 3). We first used the type of BGCs and the bacterial order to color the outer and inner circle of nodes respectively (Figure 3A). In this analysis, 31 networks of BGCs are formed, each containing between 2 to 24 nodes with 90 nodes as singletons. Each network is mainly formed by the same type of bacterial BGCs, despite the fact that a third of the networks contains one or two BGC nodes of a different type than the rest of the nodes composing this network. Additionally, in half of the networks, BGCs originated from the same bacterial order, while in the other half the BGCs originated from two or three different bacterial orders. In Figure 3B we used the species of ant and the host geography to color the inner and outer nodes respectively. However, neither the host phylogeny nor geography structures the grouping of the bacterial BGCs.

**Figure. 3.**
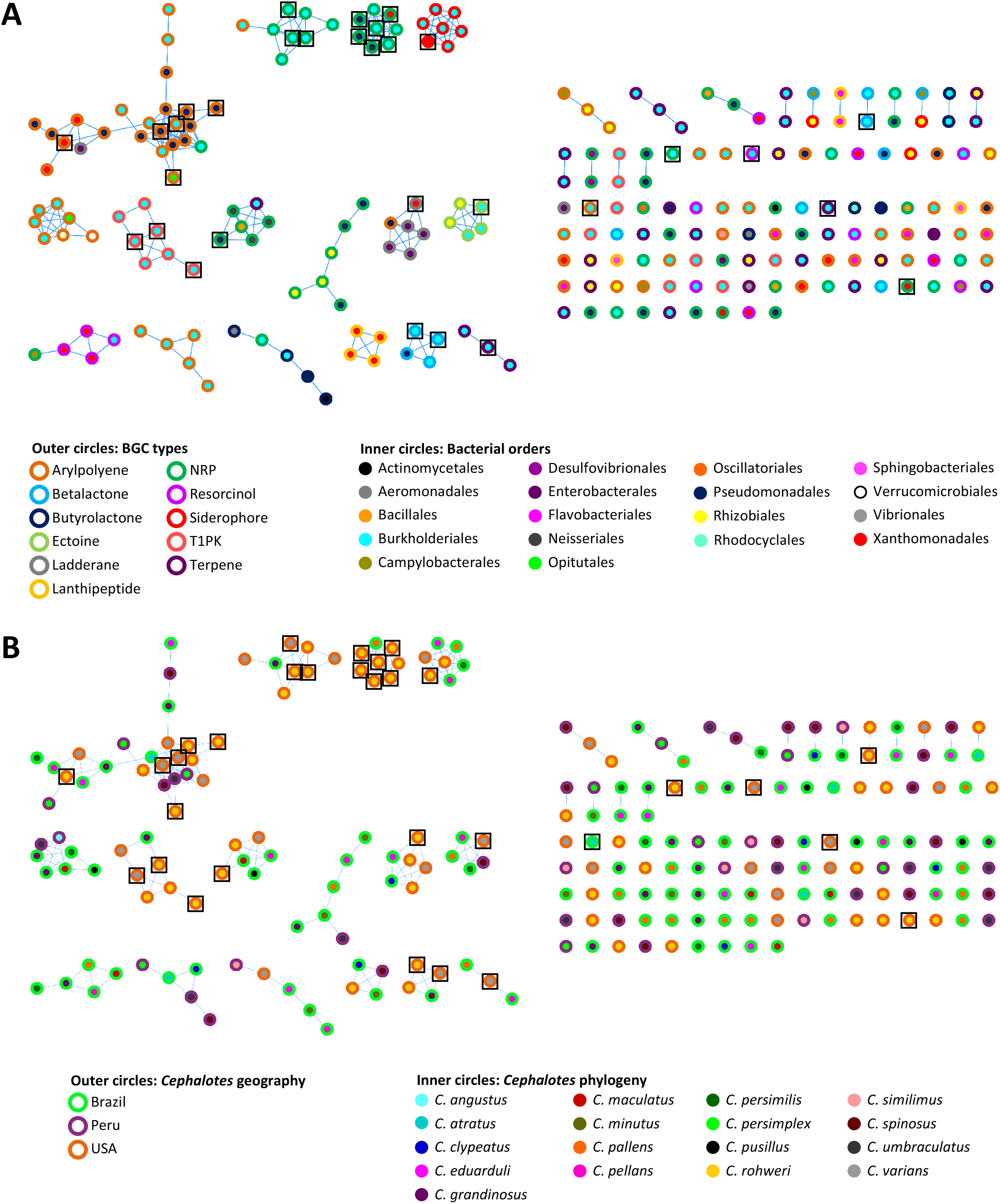
Genetic networks of the bacterial BGCs in *Cephalotes* genomes and metagenomes, with a distance threshold of 0.65 and an MCL index value of 2.0. Color codes are respective of the bacterial BGCs types and bacterial order in (A) and the *Cephalotes* geography and phylogeny in (B). Black squares represent the 31 BGCs found in bacterial genomes.

The phylogenies of the genomic core genes of the BGCs were inferred using CORASON (Figure 4, Figure S5-S6, Table S5). The four types of BGCs, arylpolyene, NRP, siderophore, and T1PK, were chosen for this analysis because they are thought to play an important role in bacteria colonization, interaction and competitiveness^51–54^. Using an arylpolyene KS gene as a query 38 different BGCs were retrieved in the arylpolyene BGC phylogeny out of the 66 detected in the *Cephalotes* bacterial genomes and metagenomes (Figure S5). These 38 arylpolyene BGCs form several distinct clades. One clade is composed of nine BGCs which are grouped in the largest arylpolyene network cluster (Figure 4A). These BGCs originated from the genomes and metagenomes of Burkholderiales and Pseudomonadales isolated from six different ant species (Figure 4A). In this clade, the arylpolyene BGCs have a length of between 24 000 to 43 000 nucleotides (nt) and between 26 to 44 genes. In the NRP BGC phylogeny based on an AMP-binding gene, 33 different NRP were retrieved out of the 57 detected in the *Cephalotes* bacterial genomes and metagenomes (Figure S6). These 33 NRP BGCs form several distinct clades including a polytomy composed of seven BGCs found in the network cluster (Figure 4B). In this clade, the NRP BGCs were identified in the genomes and the metagenome from three bacterial orders (Burkholderiales, Pseudomonadales, and Xanthomonadales) isolated from a single ant host (*C. rohweri*). The NRP BGCs have a length of between 23 000 to 53 000 nt and between 25 to 53 genes. The PKS-KS gene results in the T1PK BGC phylogeny of ten BGCs out of the 15 detected in the *Cephalotes* bacterial genomes and metagenomes (Figure 4C). All BGCs originated from the genomes and metagenomes of Burkholderiales isolated from six different ant species. The T1PK BGCs have a length of between 7 000 to 10 000 nt and between 4 to 11 genes. Seven T1PK BGCs were found a single network cluster and the three T1PK BGCs left all fall as singletons in the genetic network analysis (Figure 3A). The IucA-IucC gene was used to infer the siderophore BGC phylogeny. Eight siderophores were retrieved out of the 10 detected in the *Cephalotes* bacterial genomes and metagenomes (Figure 4D). The siderophore BGCs have a length of between 7 000 to 10 000 nt and between 4 to 11 genes. A This network matches seven siderophore BGCs from the genomes and metagenomes of Burkholderiales and Xanthomonadales each associated with a different ant species.

**Figure 4.**
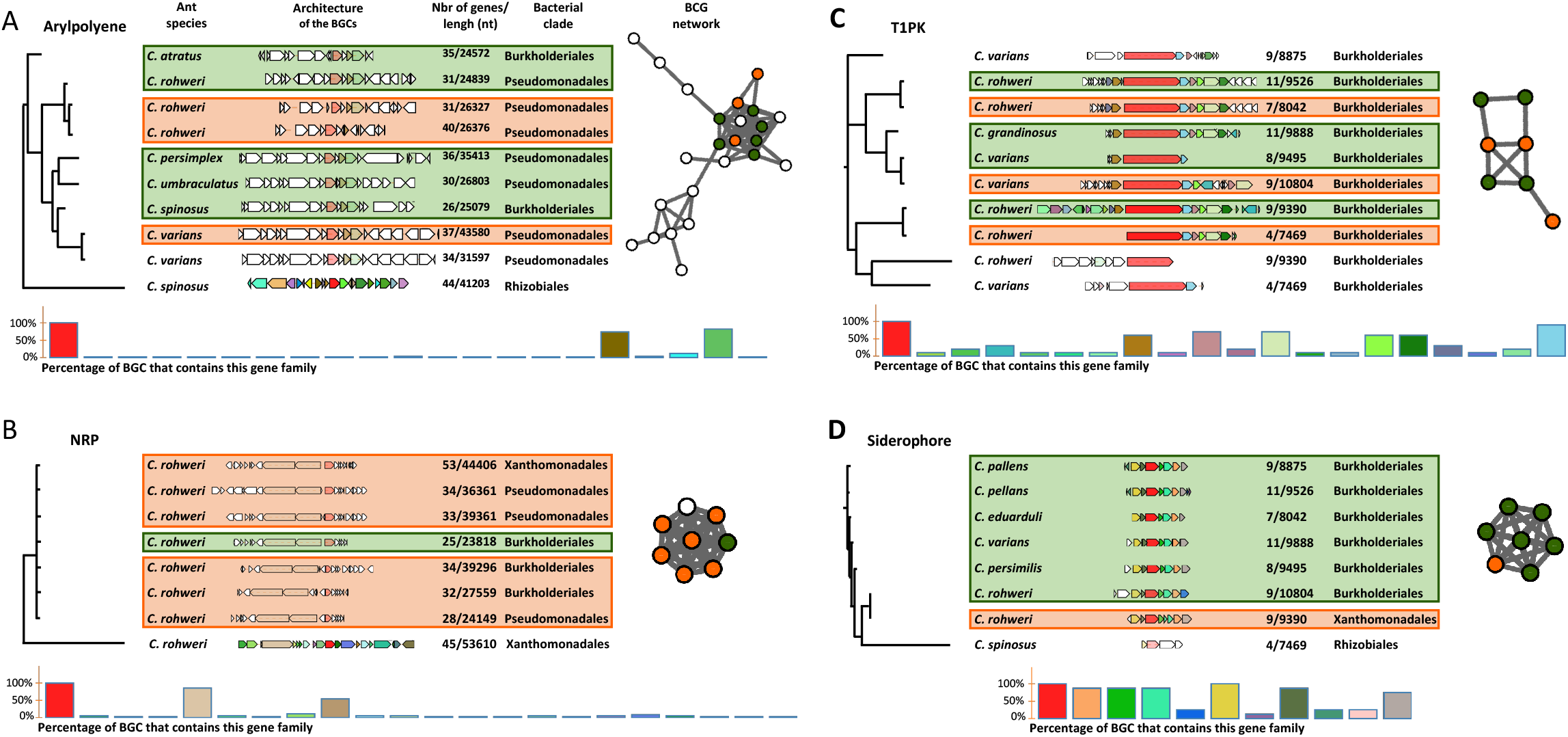
Representation of selected phylogenetic clades of arylpolyene (A), NRP (B), T1PK (C) and siderophore (D) BGCs. The bacterial origin, genomic composition, size and number of genes of each BGC, and the ant species hosting this bacterium are indicated next to the phylogeny. The histograms represent the percentage of BGC that contains this gene family. The BGCs network were retrieved from the previous analysis. Orange and green colors indicate if the BGCs originate from a bacterial genome or metagenome respectively. The white nodes in the network represent BGC which were not found in the selected phylogenetic clade. Full phylogenies of arylpolyene and NRP are found in Figures S5-S6 respectively.

The phylogenies of the elongation domains of NRP synthase and PK synthase, respectively C and KS domains, were inferred using NaPDos^35^ to check if the sequences of known metabolites were matching our data (Figure 5). The majority of the identified C domain sequences (66 out of 89) from genomes and metagenomes belong to the cyclization domain class catalyzing both peptide bond formation and subsequent cyclization. The retrieved KS domain sequences (N=13) from genomes and metagenomes fall into a single clade corresponding to *Cis*-AT modular class or the iterative domain class. Unfortunately, no close similarity with the identified C and KS sequences of known non-ribosomal peptide and polyketide were found when comparing with NaPDos and the NCBI database, thus no prediction could be made about the chemical structure of bacterial metabolites.

**Figure 5.**
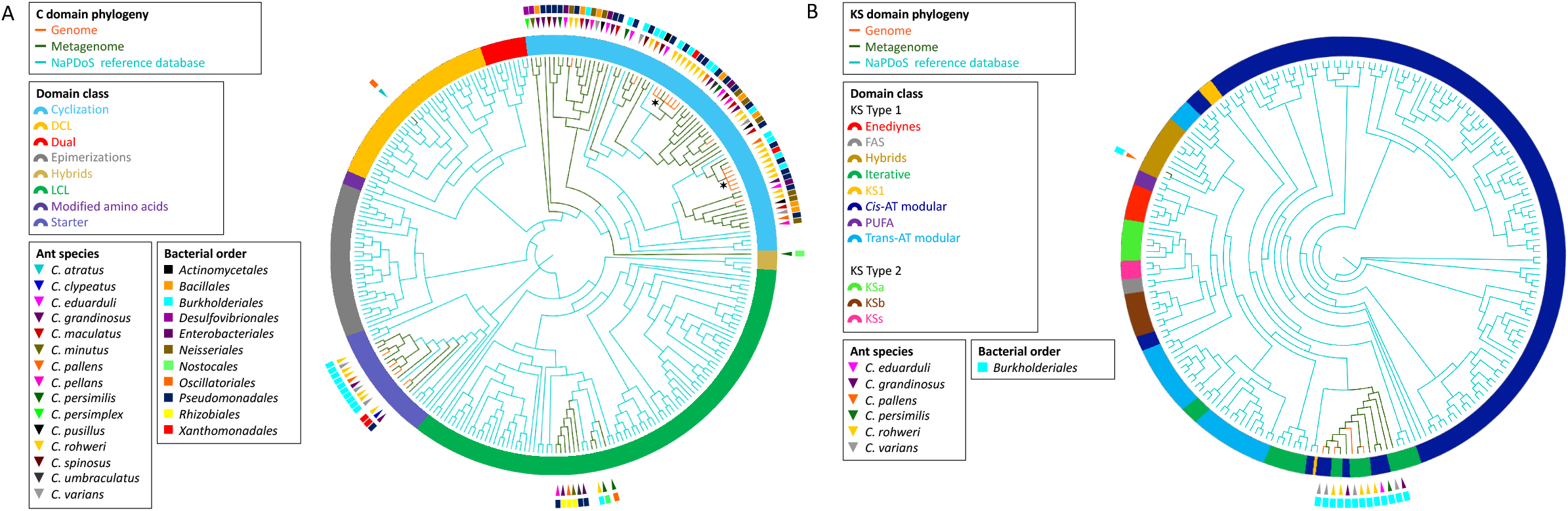
Phylogeny of retrieved C and KS elongation domain sequences in the NRP and PK assemblies respectively. The phylogenies were inferred by implementing the C domain sequences (A) and the KS domain sequences (B) from the *Cephalotes* genomes (orange branches) and metagenomes (green branches) with the NaPDoS reference database (blue branches). The ring surrounding the tips represent C domain class or KS domain class. The bacterial order from which each domain sequence was retrieved is indicated by the colored rectangle. The ant species associated with each sequence is indicated by the colored triangle. The symbols * designate the clade of the two C domain sequences find in the NRP genomic core analysis (see Fig. 4).

## Discussion

Since the multiplication of computational tools for mining microbial genomes, the discovery of BGCs of interest for pharmaceutical applications has become a growing field^19^. This approach is complementary to natural products chemistry and metabolomics in helping with the prioritization of strains synthesizing potential drugs having new chemical structures. Genome mining studies on large scale prokaryote genomic data have revealed that BGCs are distributed in large gene cluster families, with the vast majority of them still uncharacterized^41^. Some of these BGCs are distributed widely across the entire bacterial domain, with the most prominent being arylpolyenes, polyketides, saccharides and siderophores, having known and putative functions in microbe-host and microbe-microbe interactions^41^. The rise of advanced genomic tools to mine genomes and predict the chemical structure of metabolites from BGCs will certainly benefit the field of host-microbe symbiosis. Herein we present the first genome and metagenome mining study of gut bacteria across different species of herbivorous turtle ants to assess the diversity of BGCs in a nutritional symbiosis. When applied across the host phylogeny a genome mining approach provides crucial information about how the evolutionary history of the host may impact the diversity of BGCs in the symbiont genome. For instance BGCs can undergo a vast array of evolutionary process through *de novo* assembly, gene duplication/deletion, horizontal gene transfer which drive the microbial chemodiversity^55^. The systematic study of BGCs diversity of symbionts across host species from different geographic areas may provide a list of BGCs likely to play a crucial functional role for the symbiont and for the host.

Although the use of bacterial metagenomes can provide accurate genomic information^56^, one can argue that mining metagenomes using AntiSMASH^33^ may generate unreal BGCs by incorrectly assembled bins. Recently mining the metagenome for BGCs has been reported for the microbiome involved in a defensive symbiosis associated with the eggs of the beetle *Lagria villosa*^26^ and with the marine sponge *Mycale hentscheli*^57^. Our results show high similarities in the type and size of BGCs detected both in the bacterial genomes and metagenomes, with nine prominent types of BGCs detected: beta-lactone, ectoine, ladderane, NRP, resorcinol, siderophore, T1PK and terpene (Table S4 and S6). The genomic core gene analysis for arylpolyene, NRP, T1PK and siderophore show strong resemblance in the architecture and genomic core structure between bacterial genomes and metagenomes. Indeed, bacterial genomes and metagenomes of the same bacterial order and same host ant species fall together in the genetic network (Figure 3), the genomic core gene analysis (Figure 4), and the C and KS domain phylogenies (Figure 5). Together, these results show that the BGCs analysis of metagenomes are a good representation of the BGCs diversity in samples indicating that the retrieved BGCs from metagenomic data are not chimeras created by misassembling metagenomic bins.

As the 17 species of *Cephalotes* were collected in Brazil, Peru and the USA we questioned whether the host phylogeny and/or geography have an impact on symbiont BGC diversity. Our results show that the number and type of bacterial BGCs are not correlated with *Cephalotes* evolutionary history (p=0.205; Mantel test) but are more similar in *Cephalotes* samples collected in the same geographic area (p=0.004; ANOVA) (Figure 2). A significantly higher number of beta-lactone, ladderane, NRP, T1PK, and terpene BGCs associated with *Cephalotes* from the USA than associated with *Cephalotes* from South America. These results are in accordance with the geographic mosaic theory of coevolution which stipules that in strongly interacting species, geography and community ecology can shape coevolution through local adaptation^58,59^. Implications of this theory in the context of metabolites and biosynthetic gene clusters is starting to be investigated. Recently, it was shown that in soil bacteria the NRP and PK main biosynthetic domains, respectively adenylation and ketosynthase, are different across the Australian continent^60^. According to their results, the main factor for these differences is the sample latitude which can be one geographical factor explaining the differences between results from North and South America. As for the BGC architectures they are correlated with host geography (Figure 3B). This is in accordance with previous work on antibiotic-producing bacterial symbiont and fungus-growing ants^61^. In this conserved symbiosis it has been demonstrated that the geographical isolation of populations in Central America has an impact on antibiotic potency of the locally adapted symbiont but not on the BGC architecture.

Four bacterial orders possess more than 80% of all the 233 BGCs detected in the *Cephalotes* bacterial genomes and metagenomes: Burkholderiales (44%), Pseudomonadales (15%), Xanthomonadales (15%) and Rhizobiales (9%) (Figure 1, Table S4 and S6). Interestingly, these four bacterial orders belong to the *Cephalotes* core bacterial community^7^. The *Cephalotes* core bacterial community is maintained across the *Cephalotes* phylogeny and benefit these herbivorous ants with a low-nitrogen diet by recycling nitrogen from urea into essential and non-essential amino acids. In many insect nutritional symbioses the endosymbiont resides in host cells or bacteriocytes and have a reduced genome with several copies of the functional genes responsible for the biosynthesis of amino acids and vitamins to enrich in nutrients the host diet. This is contrasting with the multipartite mutualism in the gut of *Cephalotes* turtle ants. The maintenance of several bacteria is more complex compare to other symbioses involving a single gut symbiont (bean bugs^70^), or bacteriocytes endosymbionts (*Camponotus* ants^71^, beetles^9^). The conserved bacterial symbionts in *Cephalotes* have redundancy in N-metabolism and our results show these bacterial symbionts possess the vast majority of the BGCs detected. This support the idea that metabolic functions of a complex microbiome are often selected not solely for direct benefit of hosts, but for sustaining the gut community^72^. Since the *Cephalotes* core symbionts are not transmitted maternally but are newly acquired by *Cephalotes* ants in each generation, the maintenance of genes promoting gut colonization to outcompete the transient bacteria is likely high selected to be maintained.

This hypothesis is in accordance with recent findings in a bean bug which needs to acquire a specific *Burkholderia* gut symbiont from the soil environment in every new generation^70^. They demonstrated that a large number of *Burkholderia* species were able to access the bean bug gut, but in experiments of co-inoculations, the specific *Burkholderia* symbiont always outcompete all the other *Burkholderia* species. In addition in bees it has been shown that gut symbionts produce amino acids and siderophores, which are required to colonize the insect gut^72^. In the *Cephalotes* gut the five main symbionts from different bacterial orders need to colonize gut tissue, tolerate other community members and outcompete invading bacteria. The means used by co-occurring gut symbionts to survive may be through the production of specific metabolites mediating these functions. If this is true in this system, we need to identify which types of BGCs would be necessary for gut symbionts.

To test the similarities of BGC structure across different hosts, we performed CORASON genomic core gene analysis on four types of BGCs: arylpolyene, NRP, T1PK and siderophore (Figure 4, Figure S5-S6). The selected clades presented in Figure 4A and 4B show similarities in the sequence of the core genes for arylpolyene BGCs originating from Burkholderiales and Pseudomonadales, and for NRP BGCs originating from Burkholderiales, Pseudomonadales, and Xanthomonadales. These arylpolyene and NRP BGCs are each clustered in the networking analysis (Figure 3 and 4) demonstrating the architectural similarities. In the NRP BGC clade of Figure 4B the query gene AMP-binding and the two genes coding for a C domain have 100% similarity and 100% coverage across the BGCs retrieved from the six genomes and metagenomes of Burkholderiales, Pseudomonadales, and Xanthomonadales from the same host *C. rohweri*. This high similarity between the six C domain sequences from three bacterial orders is also confirmed in the phylogenetic analysis (Figure 5). This pattern suggests a possible convergence of the core genes in the gut of *C. rohweri* possibly via horizontal gene transfer. In insect-bacteria symbiosis horizontal gene transfer has been shown to occur even in bacteria going through genome reduction^26^. However the three sequences did not match any known sequence in the database limiting the identification of the bacterial origin of these genes. On the other hand, the T1PK and siderophore core gene phylogenies (Figure 4B and 4C) show that BGCs from Burkholderiales across different host species share a strong likely genetic conservation. Several identical genes of T1PK and siderophore BGCs are present across the core gene phylogenies supporting a conserved architecture which suggests a common origin or convergent evolution of Burkholderiales BGCs across *Cephalotes* species. Symbiotic convergent evolution has been reported in insect-bacteria symbiosis^74,75^, showing insect symbionts retain the ability to produce a certain number of proteins and functions which have an importance for both bacteria and host survival.

A final remark concerns the potential division of labor in a multipartite mutualism. In honey bees the gut microbiota are dominated by five coevolved bacterial clusters. However the gut bacterial species have different abilities in digesting polysaccharides which can be explained by a divergence into different ecological niches inside the gut of their hosts^8^. In the *Cephalotes* gut four of the five conserved symbionts possess the majority of the retrieved BGCs but Opitutales a major player in the *Cephalotes* nutritional symbiosis is lacking BGCs. Although multiple bins of Opitutales were retrieved only two BGCs (arylpolyene) were detected (Figure S7). In the *Cephalotes* gut only Opitutales possess urease genes able to transform urea into ammonia readily used by the four other symbionts. Opitutales are localized in the midgut whereas Burkholderiales, Pseudomonadales, Rhizobiales, and Xanthomonadales are found in the ileum of the hindgut (Flynn et al. in prep.). This spatial isolation prevents direct interaction of Opitutales with other symbionts and their lack of BCGs supports the idea that BGCs have an essential role for symbionts living in close proximity.

Our study sheds light on the question of why many obligate host-associated, non-endosymbiotic bacteria do not have highly reduced genomes, like their endosymbiotic counterparts. In this multipartite nutritional symbiosis with herbivorous turtle ants the core bacterial community members not only maintain genes to supplement the host’s nitrogen-poor diet^7^, but we show they also retain BGCs that likely contribute to their colonization of the gut and mediate bacterial-bacterial communication and interactions. Future work on host-associated bacterial communities should begin to explore the specific metabolites expressed by these microbes and their functional outcome in these complex ecosystems.

## Author Contributions

AC, CSM, and CD conceived the study. AC performed the analyses. AC, CSM, and CD interpreted the results. AC drafted the manuscript. AC, CSM, and CD revised the manuscript.

## Funding

This work received financial support from an “Investissement d’Avenir” grant managed by Agence Nationale de la Recherche (CEBA, ref. ANR-10-LABX-25-01) to CD, a National Science Foundation (NSF DEB 1900357) to CSM. We are grateful to the University of French Guiana for Anaïs Chanson’s Ph.D. fellowship and traveling grants.

## Conflict of Interest

The authors declare no competing financial and non-financial competing interests.

## Acknowledgments

The authors are grateful to Yi Hu and Jon Sanders for their knowledge and advice on bacterial metagenomes analysis and to Manuela Ramalho for her unfailing assistance with statistical analysis. The authors are thankful to Simon Shaw for his help with the detection of metagenomic biosynthetic gene clusters, and to Nelly Selem for her assistance with the genomic core analysis.

## Notes

### Competing Interest Statement

The authors have declared no competing interest.

