## Supplementary material for "Assessing biosynthetic gene cluster diversity in a multipartite nutritional symbiosis between herbivorous turtle ants and conserved gut symbionts": SI

### Table of contents

Figure S1. Summary of the *Cephalotes* genomes and metagenomes workflow. 2

Figure S2. Assessment of the differences between the BGCs in the genome analysis. 3

Figure S3. Assessment of the metagenomic binning method quality. 4

Figure S4. Assessment of the bacterial composition of the metagenomic bins. 5

Figure S5. Genomic core analysis and phylogenetic tree of the arylpolyene BGCs from the Cephalotes bacterial genomes and metagenomes. 6

Figure S6. Genomic core analysis and phylogenetic tree of the NRP BGCs from the Cephalotes bacterial genomes and metagenomes. 7

Figure S7. Number and type of BGCs identified in the genomes and metagenomes of the *Cephalotes* gut core bacterium. 8

Table S1. Assembly statistics of genomes. 9

Table S2. Assembly statistics of metagenomes. 10

Table S3. Reference BGC and query protein selected in the genomic core analysis. 11

Table S4. BGCs types identified in the bacterial genomes 12

Table S5. Statistic assessment of the metagenomic bins created. 13

Table S6. BGCs types identified in the bacterial metagenomes. 14

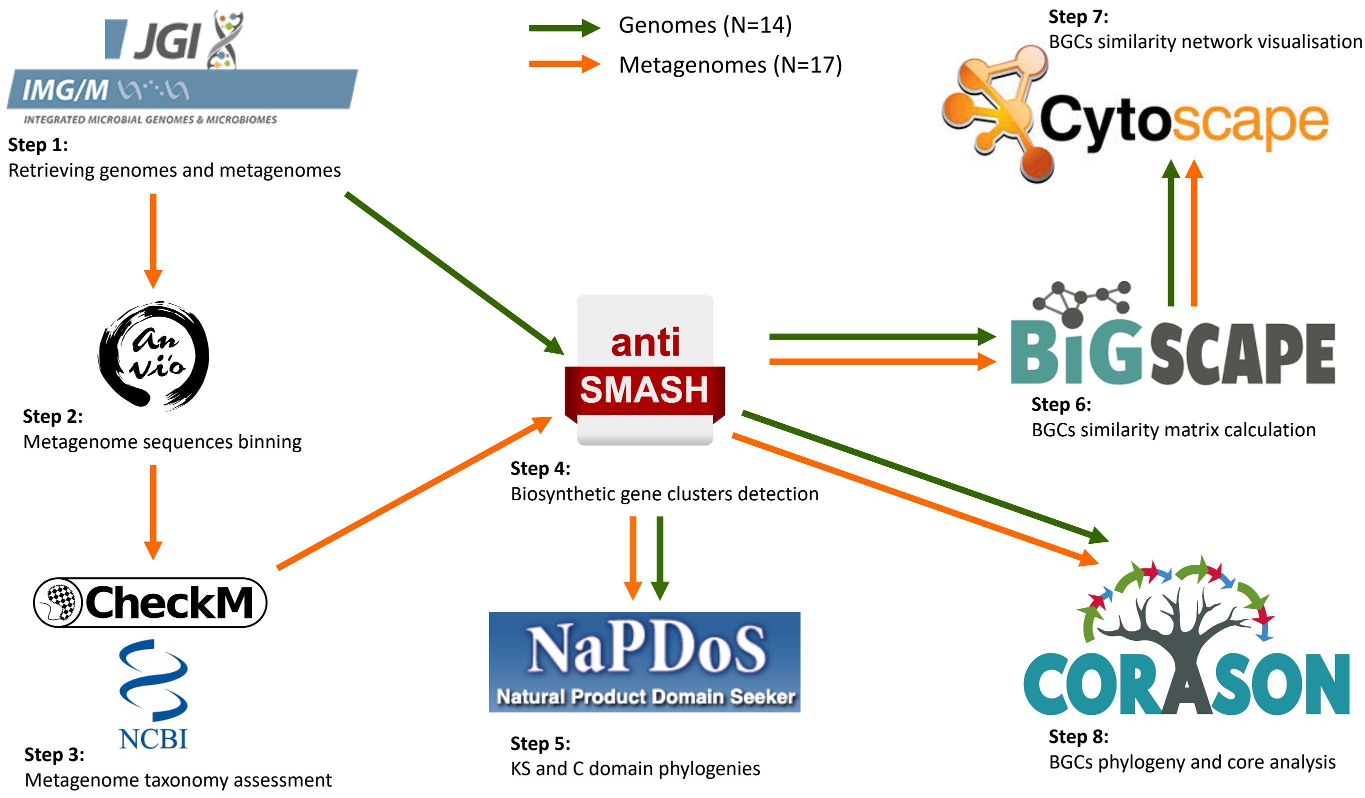


**Figure S1. Summary of the *Cephalotes* genomes and metagenomes workflow.**

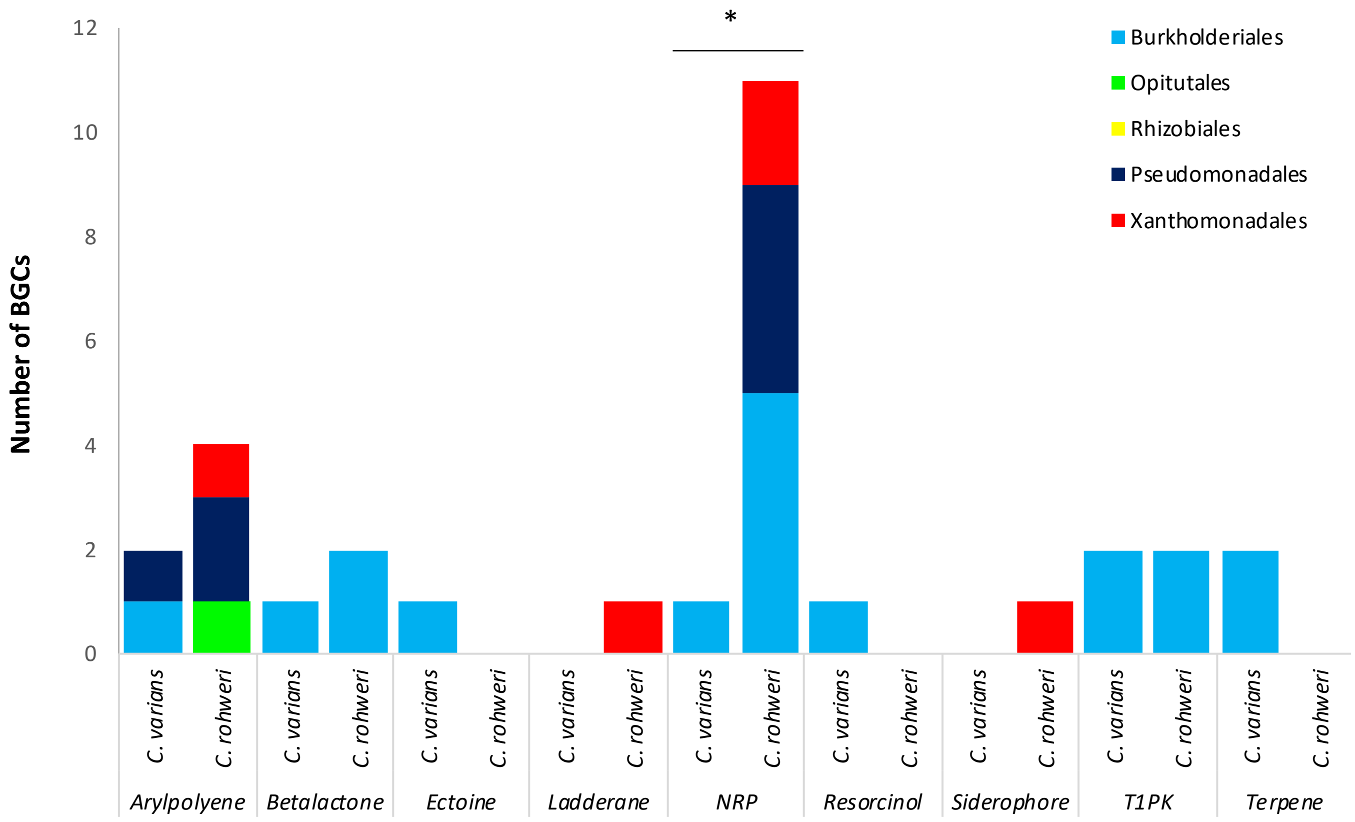


**Figure S2. Assessment of the differences between the BGCs in the genome analysis**.
One-way Anova showing the statistical differences between the bacterial BGCs across bacteria species and Cephalotes species. The symbol * represents a p-value lower than 0.05.

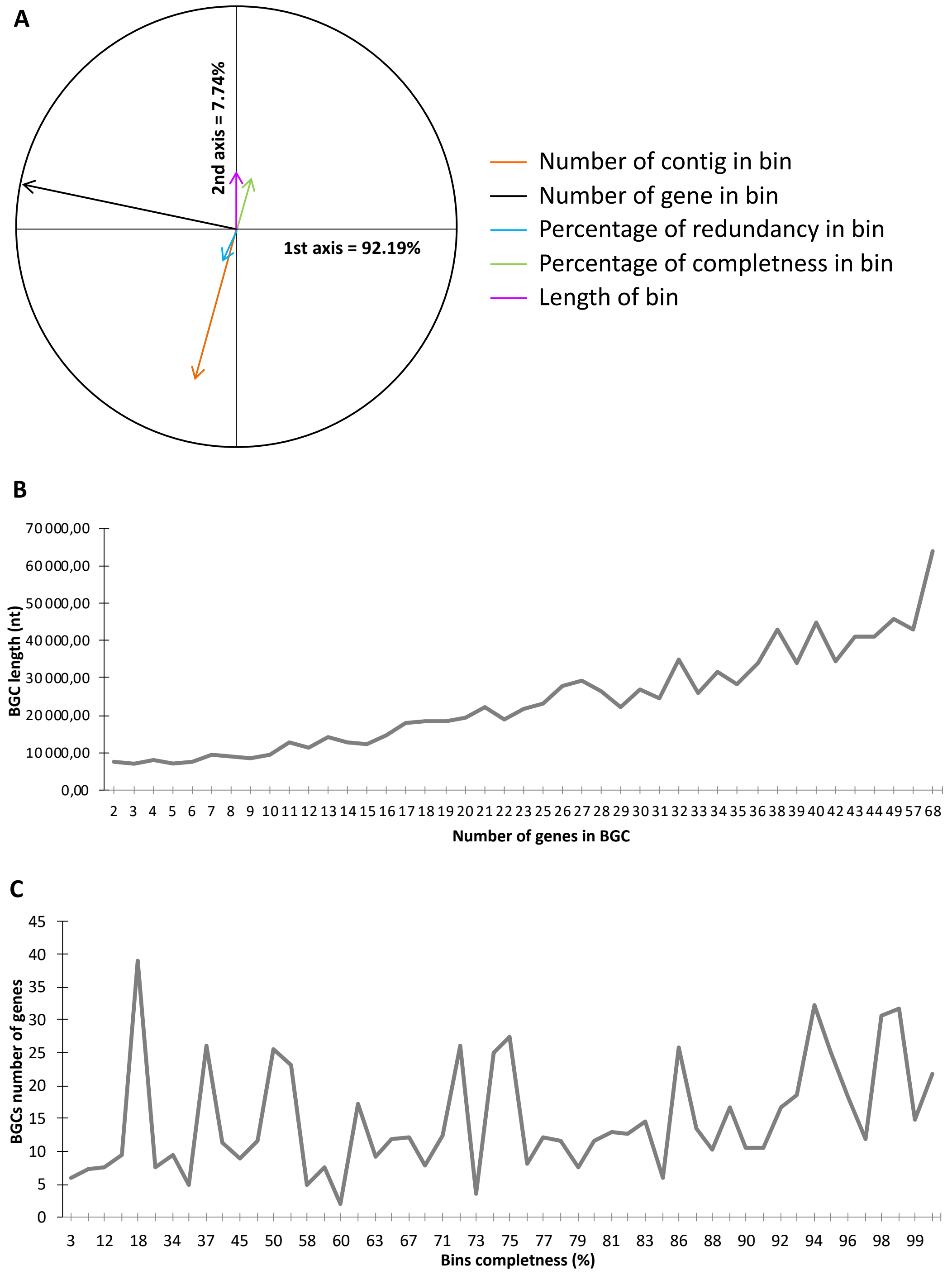


**Figure S3. Assessment of the metagenomic binning method quality.**

(A) Variable correlation plots generated for the principal component analysis plot of bin completeness (N=168). (B) Spearman correlation between the length and the number of gene in each BGC. (N=233; S=354,810; p<2.2e-16; rho=0.832) (C) Spearman correlation between the length of BGC and the completeness of the bin in which the BGC was found (N=233; S=1,579,300; p=9.252e-13; rho=0.251).

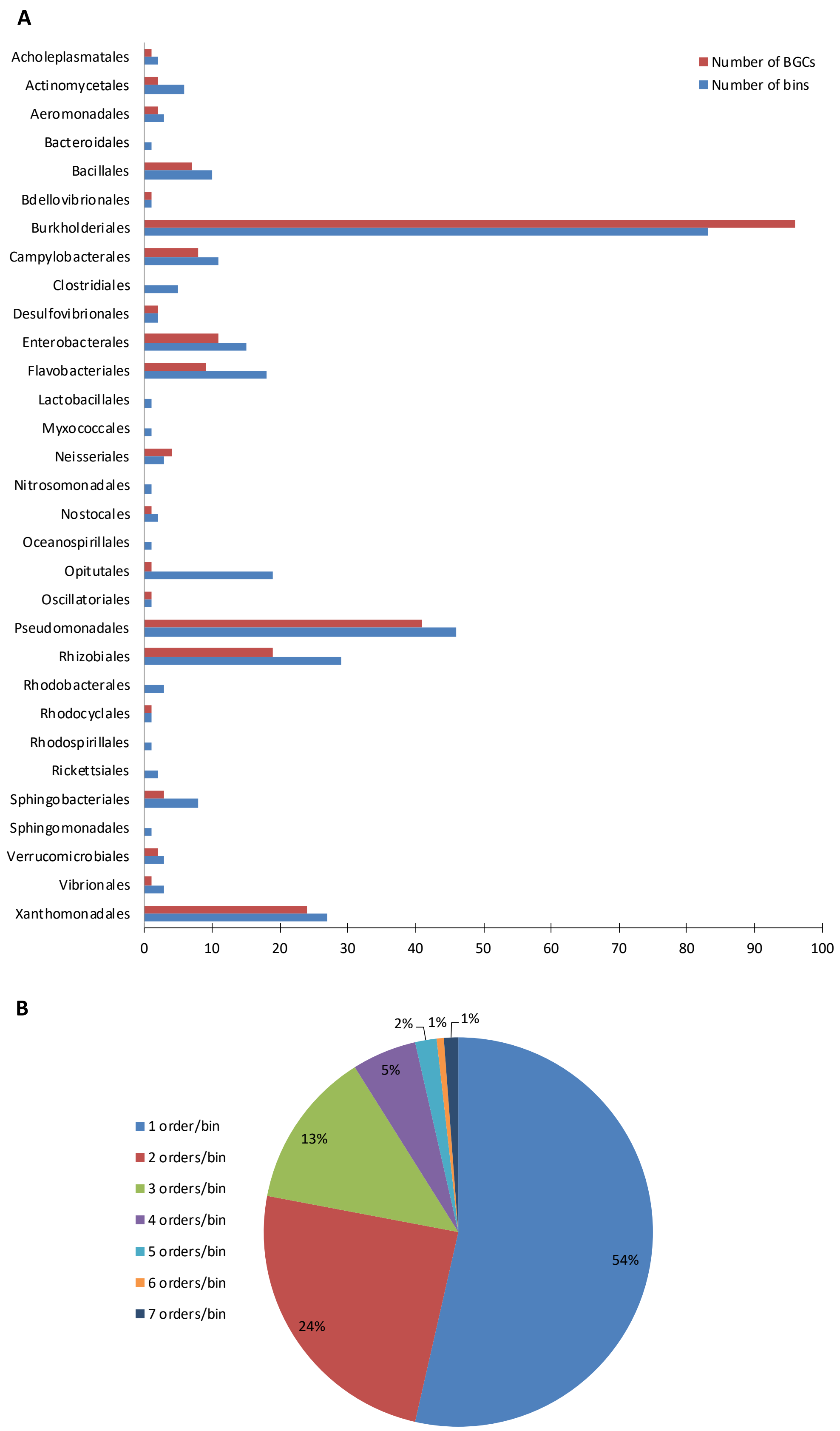


**Figure S4. Assessment of the bacterial composition of the metagenomic bins.**
(A) List of bacterial orders identified in the bins identified in metagenomes. The blue bars represent the number of bins in which each bacterial order was found. The red bars represent the number of BGCs possessed by each bacterial order. (B) The proportion of different bacterial orders found in the bin.

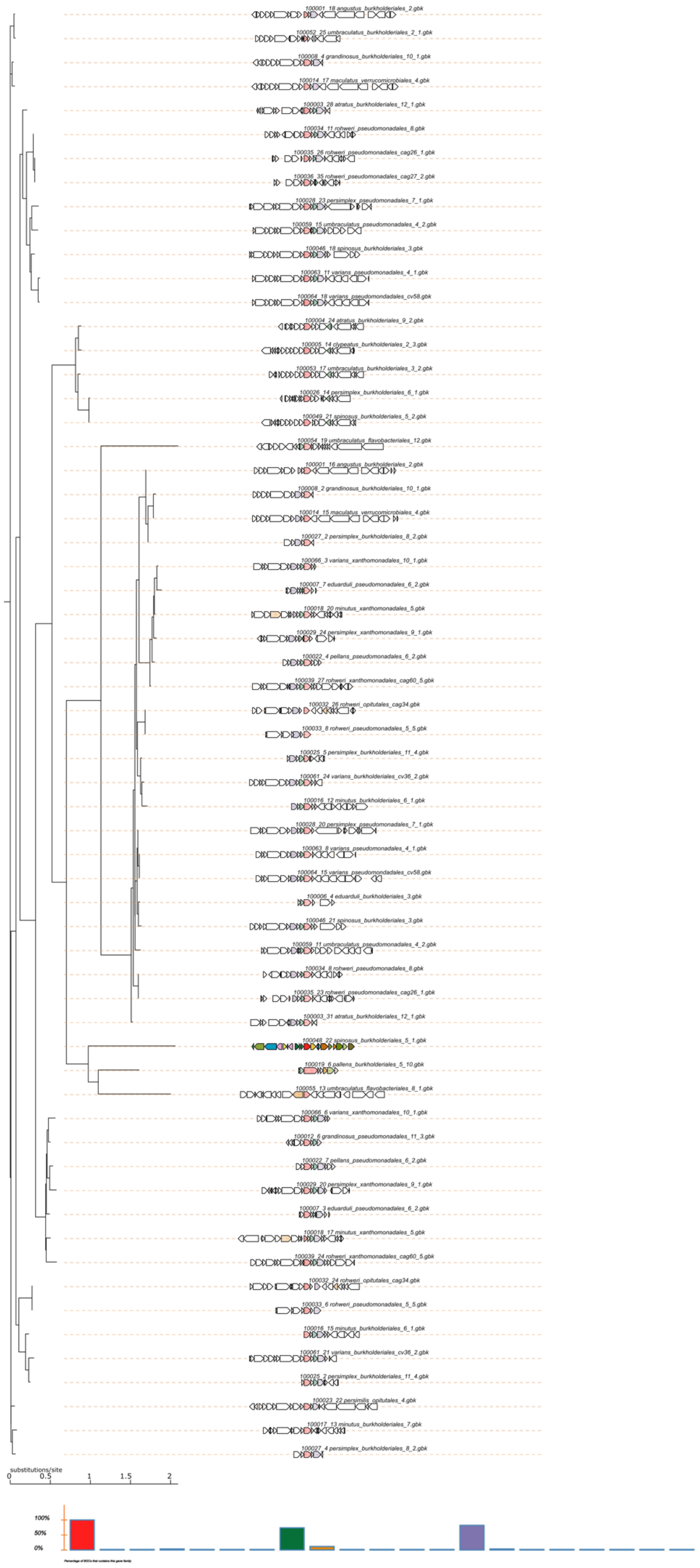

**Figure S5. Genomic core analysis and phylogenetic tree of the arylpolyene BGCs from the Cephalotes bacterial genomes and metagenomes.**

This phylogeny was implemented through CORASON by uploading all the arylpolyene BCGs retrieved from the *Cephalotes* associated genomes and metagenomes into a single database and by comparing their sequences and architectures to a reference BGC and query protein as detailed in Table S3. The histogram represents the percentage of BGC that contains this gene family.

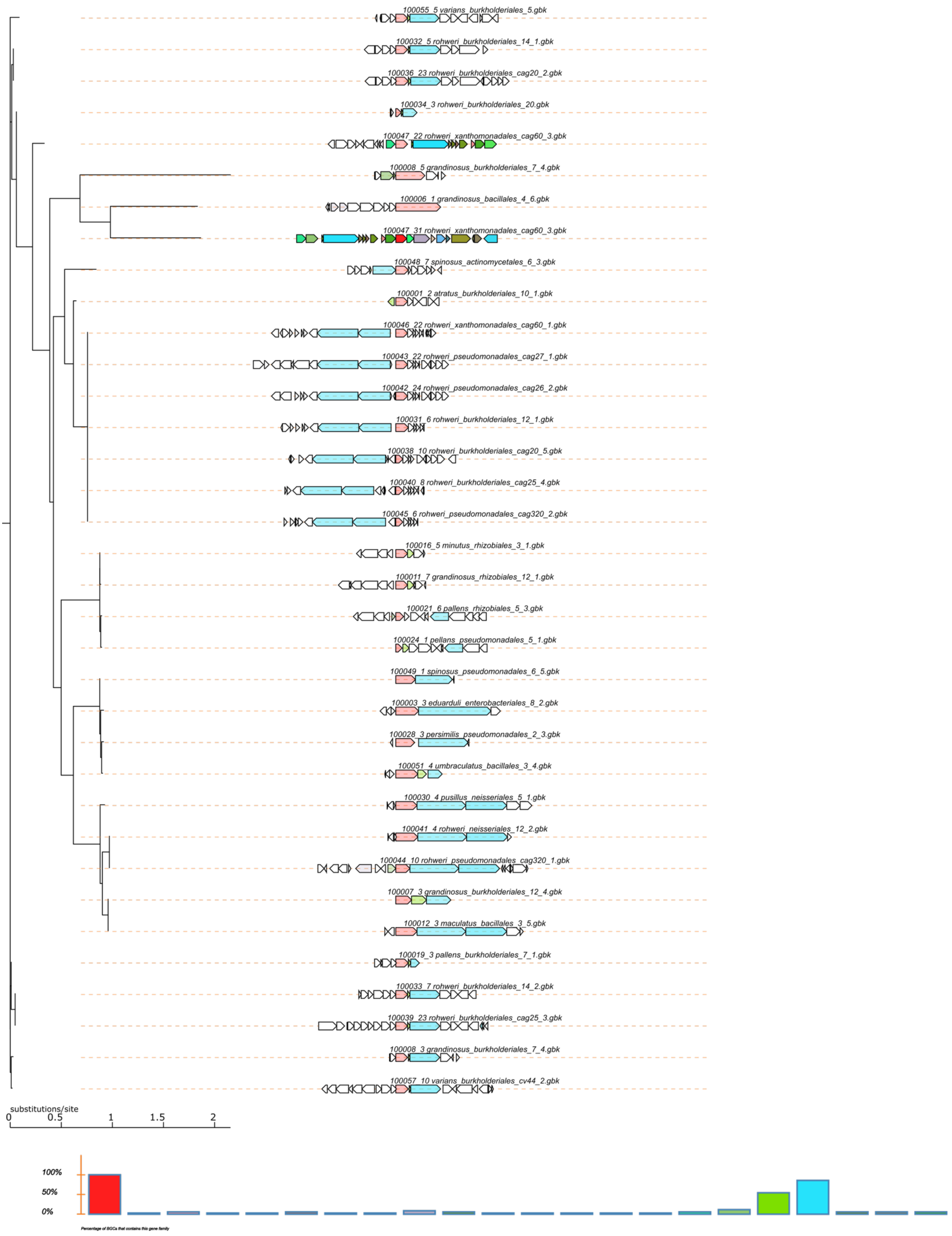


**Figure S6. Genomic core analysis and phylogenetic tree of the NRP BGCs from the Cephalotes bacterial genomes and metagenomes.**

This phylogeny was implemented through CORASON by uploading all the NRP BCGs retrieved from the *Cephalotes* associated genomes and metagenomes into a single database and comparing their sequences and architectures to a reference BGC and query protein as detailed in Table S3. The histogram represents the percentage of BGC that contains this gene family.

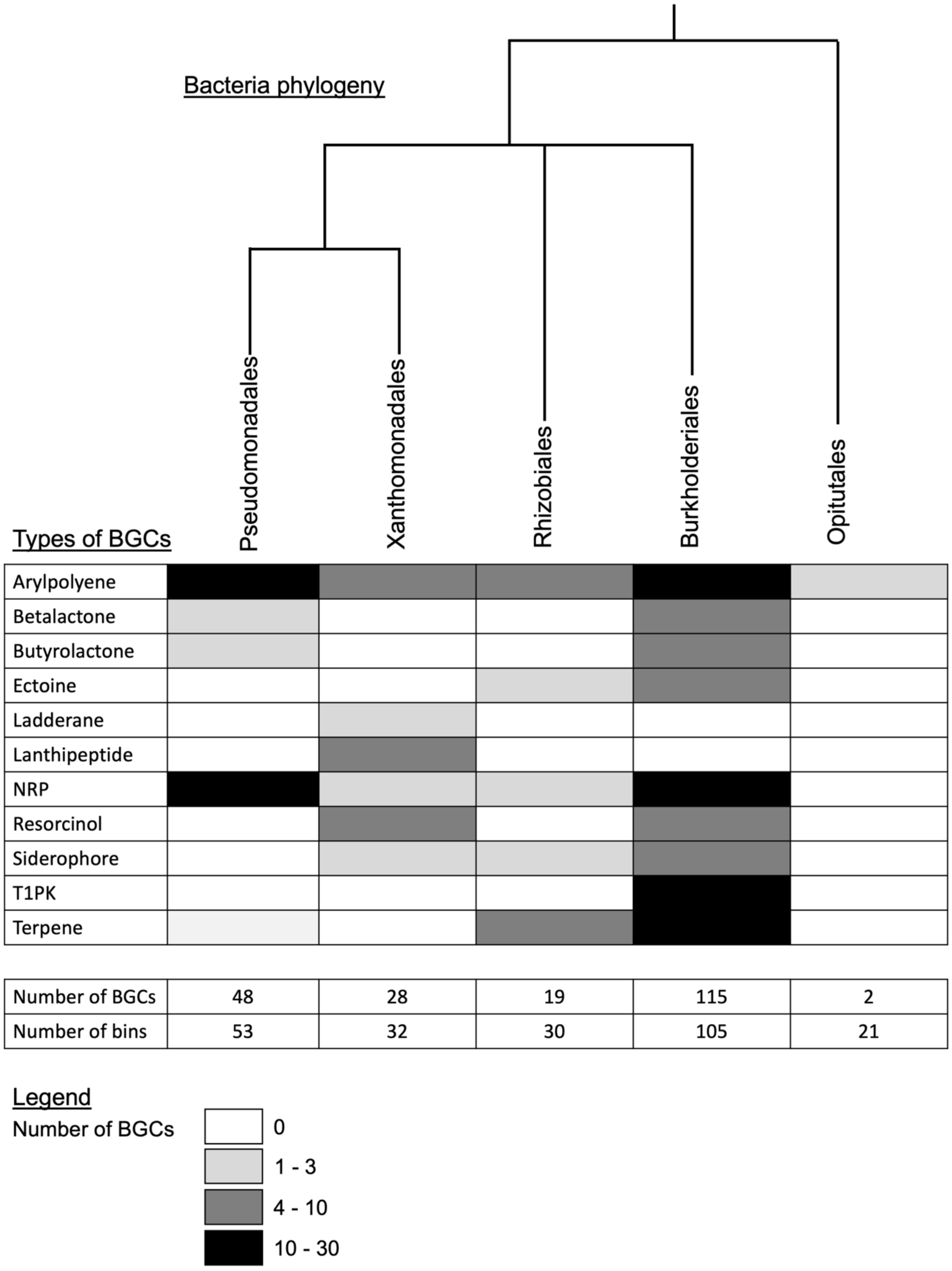


**Figure S7. Number and type of BGCs identified in the genomes and metagenomes of the *Cephalotes* gut core bacterium.**

The black to white shading filling in the table represent the number of BGCs of each type identified in the genomes and metagenomes of the 5 main bacterial order associated with the *Cephalotes* gut. The blue shading filling represent the number of bins in which each bacterial order has been found.

**Table S1. Assembly statistics of genomes.**
This table was extracted from the Supplementary Information of Hu *et al*., 2018.

| Genome name | Total length (bp) | Scaffold number | N50 (bp) | GC% mean | IMG project ID | Ant host |
|---|---|---|---|---|---|---|
| *Cephaloticoccus primus* Cag34 | 2,353,466 | 23 | 207,028 | 62.43% | Gp0154034 | *C. rohweri* |
| *Cephaloticoccus canophilus* Cv41 | 2,094,663 | 34 | 169,871 | 59.29% | Gp0110136 | *C. varians* |
| Rhizobiales *sp.* JR021-5 | 1,943,462 | 1 | 1,943,462 | 57.28% | Gp0155340 | *C. varians* |
| Xanthomonadales *sp.* Cag60 | 3,424,135 | 179 | 44,406 | 56.17% | Gp0127963 | *C. rohweri* |
| *Ventosimonas gracilis* Cv58 | 2,623,100 | 221 | 58,779 | 53.40% | Gp0110146 | *C. varians* |
| *Ventosimonas sp.* Cag26 | 2,729,219 | 92 | 79,155 | 54.82% | Gp0154031 | *C. rohweri* |
| *Ventosimonas sp.* Cag27 | 2,719,975 | 89 | 79,155 | 54.89% | Gp0154032 | *C. rohweri* |
| *Ventosimonas sp.* Cag320 | 2,825,500 | 94 | 108,424 | 55.67% | Gp0154033 | *C. rohweri* |
| Burkholderiales *sp.* Cag20 | 3,101,960 | 121 | 69,296 | 58.26% | Gp0154021 | *C. rohweri* |
| Burkholderiales *sp.* Cag25 | 3,191,349 | 112 | 142,682 | 58.13% | Gp0154030 | *C. rohweri* |
| Burkholderiales *sp.* Cv44 | 3,159,338 | 601 | 33,742 | 60.40% | Gp0110143 | *C. varians* |
| Burkholderiales *sp.* Cv33a | 2,469,676 | 150 | 48,592 | 58.91% | Gp0110144 | *C. varians* |
| Burkholderiales *sp.* Cv26 | 2,921,795 | 234 | 54,338 | 59.95% | Gp0110137 | *C. varians* |
| Burkholderiales *sp.* Cv52 | 2,935,707 | 203 | 93,689 | 60.36% | Gp0110145 | *C. varians* |

**Table S2. Assembly statistics of metagenomes.**

| Ant species | Reads numbers | Total length Mbp | Total >=1 Kbp scaffold length (Mbp) | Scaffold number | Scaffold number (>=1Kbp) | N50 for scaffold length (bp) | GC% mean | Read coverage mean | IMG project ID |
|---|---|---|---|---|---|---|---|---|---|
| *C. varians* PL005W | 40,804,316 | 184.98 | 142.43 | 281,884 | 129,829 | 1,074 | 39.18% | 16.60 | Gp0095985 |
| *C. varians* PL010W* | 135,924,092 | 436.39 | 340.3 | 647,079 | 91,083 | 10,750 | 41.95% | 41.75 | Gp009598 |
| *C. angustus* | 3,534,316 | 21.5 | 8.22 | 30,858 | 3,531 | 1,118 | 50.80% | 6.79 | Gp0125961 |
| *C. atratus* | 34,205,408 | 218.88 | 141.37 | 198,281 | 67,767 | 1,488 | 38.53% | 9.20 | Gp0125962 |
| *C. clypeatus* | 24,093,876 | 104.18 | 41.7 | 135,743 | 16,952 | 1,031 | 37.42% | 8.62 | Gp0125963 |
| *C. eduarduli* | 22,505,270 | 82.63 | 37.38 | 100,267 | 12,793 | 1,259 | 39.99% | 9.53 | Gp0125964 |
| *C. grandinosus* | 37,871,910 | 156.24 | 60.81 | 220,221 | 22,090 | 1,215 | 39.68% | 10.06 | Gp0125967 |
| *C. maculatus* | 40,739,970 | 144.43 | 42.17 | 230,315 | 18,066 | 921 | 39.60% | 7.52 | Gp0125968 |
| *C. minutus* | 31,576,424 | 115.64 | 37.69 | 180,629 | 15,101 | 1,038 | 38.38% | 8.10 | Gp0125969 |
| *C. pallens* | 28,073,250 | 148.44 | 55.05 | 211,667 | 21,214 | 1,095 | 39.56% | 7.92 | Gp0125970 |
| *C. pellans* | 33,836,572 | 142.82 | 43.13 | 224,021 | 18,268 | 934 | 38.32% | 8.51 | Gp0126569 |
| *C. persimilis* | 28,044,242 | 106.24 | 44.84 | 133,984 | 19,037 | 1,077 | 40.41% | 10.09 | Gp0126571 |
| *C. persimplex* | 40,440,804 | 231.49 | 102.43 | 291,903 | 53,617 | 1,090 | 37.94% | 10.14 | Gp0126580 |
| *C. pusillus* | 18,252,188 | 61.89 | 24.03 | 93,263 | 9,713 | 1,389 | 42.35% | 6.67 | Gp0126572 |
| *C. rohweri* | 58,943,942 | 305.84 | 271.58 | 142,961 | 78,950 | 3,982 | 40.47% | 22.40 | Gp0126573 |
| *C. similimus* | 30,304,070 | 92.55 | 37.04 | 122,379 | 16,294 | 1,030 | 37.75% | 13.09 | Gp0126574 |
| *C. spinosus* | 19,456,806 | 94.81 | 41 | 117,242 | 13,728 | 1,206 | 38.79% | 8.66 | Gp0126575 |
| *C. umbraculatus* | 37,344,984 | 256.42 | 193.69 | 191,339 | 84,591 | 1,984 | 37.76% | 14.53 | Gp0126577 |

*This metagenome was excluded in our analysis

This table was extracted from the Supplementary Information of :

Hu, Y. *et al.* Herbivorous turtle ants obtain essential nutrients from a highly conserved nitrogen-recycling gut microbiome. *Nat. Commun.* (2018).

**Table S3. Reference BGC and query protein selected in the genomic core analysis.**

| Database | Reference BGC | Reference BGC length (bp) | Reference BGC length (gene number) | Query protein domain | Query protein coordinates in the reference BGC |
|---|---|---|---|---|---|
| Arylpolyene | *C. spinosus* Bin_5_1 | 41,203 | 44 | APE_KS2 | 38,905-40,107 |
| NRP | *C. rohweri* Xanthomonadales Cag60_3 | 53,609 | 45 | AMP-binding | 64,471-65,991 |
| T1PK | *C. rohweri* Burkholderiales Cag25_2 | 47,911 | 35 | PKS_KS | 197,260-205,170 |
| Siderophore | *C. pellans* Bin_2_7 | 9,526 | 11 | IucA_IucC | 2,926-4,788 |

**Table S4. BGCs types identified in the bacterial genomes**

| Host | Symbiont | Type of cluster | Number of genes | Size of cluster (nt) |
|---|---|---|---|---|
| *C. varians* | Burkholderiales Cv33a | | | |
| *C. varians* | Burkholderiales Cv36_1 | Terpene | 20 | 21,712 |
| *C. varians* | Burkholderiales Cv36_2 | Arylpoyene | 27 | 25,774 |
| *C. varians* | Burkholderiales Cv44_1 | T1PKS | 42 | 47,950 |
| *C. varians* | Burkholderiales Cv44_2 | NRPS | 27 | 33,193 |
| *C. varians* | Burkholderiales Cv44_3 | Betalactone | 16 | 15,437 |
| *C. varians* | Burkholderiales Cv52_1 | Terpene | 22 | 21,745 |
| *C. varians* | Burkholderiales Cv52_2 | Resorcinol | 31 | 41,911 |
| *C. varians* | Burkholderiales Cv52_3 | T1PKS | 28 | 34,252 |
| *C. varians* | Burkholderiales Cv52_4 | Ectoine | 8 | 9,669 |
| *C. varians* | *Cephaloticoccus canophilus* Cv41 | | | |
| *C. varians* | Rhizobiales JR021-5 | | | |
| *C. varians* | *Ventosimonas gracilis* Cv58 | Arylpoyene | 37 | 43,580 |
| *C. rohweri* | Burkholderiales Cag20_1 | T1PKS | 36 | 36,938 |
| *C. rohweri* | Burkholderiales Cag20_2 | NRPS | 27 | 30,210 |
| *C. rohweri* | Burkholderiales Cag20_3 | Betalactone | 20 | 22,669 |
| *C. rohweri* | Burkholderiales Cag20_4 | NRPS | 7 | 9,722 |
| *C. rohweri* | Burkholderiales Cag20_5 | NRPS | 34 | 39,296 |
| *C. rohweri* | Burkholderiales Cag25_1 | Betalactone | 20 | 28,119 |
| *C. rohweri* | Burkholderiales Cag25_2 | T1PKS | 34 | 47,910 |
| *C. rohweri* | Burkholderiales Cag25_3 | NRPS | 41 | 45,968 |
| *C. rohweri* | Burkholderiales Cag25_4 | NRPS | 32 | 27,559 |
| *C. rohweri* | *Cephaloticoccus primus* Cag34 | Arylpoyene | 47 | 42,932 |
| *C. rohweri* | *Ventosimonas* Cag26_1 | Arylpoyene | 31 | 26,327 |
| *C. rohweri* | *Ventosimonas* Cag26_2 | NRPS | 33 | 39,361 |
| *C. rohweri* | *Ventosimonas* Cag27_1 | NRPS | 34 | 36,361 |
| *C. rohweri* | *Ventosimonas* Cag27_2 | Arylpoyene | 40 | 26,376 |
| *C. rohweri* | *Ventosimonas* Cag320_1 | NRPS | 18 | 28,447 |
| *C. rohweri* | *Ventosimonas* Cag320_2 | NRPS | 28 | 24,149 |
| *C. rohweri* | Xanthomonadales Cag60_1 | NRPS | 53 | 44,406 |
| *C. rohweri* | Xanthomonadales Cag60_2 | Ladderane | 28 | 29,577 |
| *C. rohweri* | Xanthomonadales Cag60_3 | NRPS | 45 | 53,610 |
| *C. rohweri* | Xanthomonadales Cag60_4 | Siderophore | 9 | 9,390 |
| *C. rohweri* | Xanthomonadales Cag60_5 | Arylpoyene | 43 | 38,815 |

This table contains the number of BGCs detected in each bacteria isolated from *C. varians* and *C.* rohweri, as well as the length and number of genes composing these BGCs. The colors fill of the cell represent the type of BGCs.

**Table S5. Statistic assessment of the metagenomic bins created.**

Additional file

Table_S5_List_metagenomes_bin.xlsx

The metagenomic bins were created by uploading each metagenome sequences into the Anvi'o version 5.5. This table contains for each *Cephalotes* metagenome the number of bins created, and for each bin it shows the percentage of completeness, the percentage of redundancy, the number of contigs, the number of genes, and the number of BGCs found. All these data were obtained through the Anvi'o software.

**Table S6. BGCs types identified in the bacterial metagenomes.**

Additional file

Table_S6_List_metagenomes_BGC.xlsx

Types and origins of the bacterial BGCs in the metagenome analysis. The colors of the cell represent the type of BGCs. The colors of police of the symbionts represent the bacterial order. The code color is the same than in Fig. 2.
