## Supplementary material for "Assessing biosynthetic gene cluster diversity in a multipartite nutritional symbiosis between herbivorous turtle ants and conserved gut symbionts": Table S5

| Host species | Number of bins | | Completeness bin (%) | Redundancy bin (%) |
|---|---|---|---|---|
| C_angustus | 2 | 1 | 17.99 | 5.76 |
| | | 2 | 97.12 | 0.00 |
| C_atratus | 15 | 1 | 95.68 | 5.76 |
| | | 2 | 97.12 | 3.60 |
| | | 3 | 74.10 | 38.13 |
| | | 4 | 97.12 | 0.72 |
| | | 5 | 99.28 | 0.00 |
| | | 6 | 87.05 | 20.14 |
| | | 7 | 82.73 | 1.44 |
| | | 8 | 97.12 | 1.44 |
| | | 9 | 80.58 | 24.46 |
| | | 10 | 79.14 | 41.73 |
| | | 11 | 76.98 | 25.90 |
| | | 12 | 64.03 | 35.25 |
| | | 13 | 94.96 | 1.44 |
| | | 14 | 82.01 | 3.60 |
| | | 15 | 3.60 | 0.00 |
| C_clypeatus | 6 | 1 | 98.56 | 56.83 |
| | | 2 | 98.56 | 29.50 |
| | | 3 | 97.12 | 52.52 |
| | | 4 | 99.28 | 0.00 |
| | | 5 | 73.38 | 0.72 |
| | | 6 | 97.12 | 4.32 |
| C_eduarduli | 8 | 1 | 97.12 | 0.00 |
| | | 2 | 88.49 | 3.60 |
| | | 3 | 99.28 | 6.47 |
| | | 4 | 89.21 | 10.07 |
| | | 5 | 97.12 | 0.00 |
| | | 6 | 82.01 | 17.27 |
| | | 7 | 72.66 | 12.95 |
| | | 8 | 18.71 | 3.60 |
| C_grandinosus | 12 | 1 | 97.84 | 0.72 |
| | | 2 | 76.98 | 2.16 |
| | | 3 | 99.28 | 1.44 |
| | | 4 | 79.14 | 24.46 |
| | | 5 | 81.29 | 36.69 |
| | | 6 | 58.27 | 29.50 |
| | | 7 | 58.27 | 28.06 |
| | | 8 | 42.45 | 12.23 |
| | | 9 | 58.27 | 34.53 |
| | | 10 | 67.63 | 46.04 |
| | | 11 | 66.91 | 15.83 |
| | | 12 | 45.32 | 19.42 |
| C_maculatus | 6 | 1 | 99.28 | 2.16 |
| | | 2 | 99.28 | 1.44 |
| | | 3 | 63.31 | 28.78 |
| | | 4 | 95.68 | 0.00 |
| | | 5 | 60.43 | 18.71 |
| | | 6 | 93.53 | 5.04 |
| C_minutus | 8 | 1 | 94.96 | 0.72 |

| | | | | |
|---|---|---|---|---|
| | | 2 | 97.12 | 1.44 |
| | | 3 | 7.19 | 0.00 |
| | | 4 | 76.98 | 2.16 |
| | | 5 | 98.56 | 0.72 |
| | | 6 | 71.94 | 3.60 |
| | | 7 | 97.84 | 1.44 |
| | | 8 | 77.70 | 38.85 |
| C_pallens | 10 | 1 | 95.68 | 0.72 |
| | | 2 | 93.53 | 1.44 |
| | | 3 | 98.56 | 6.47 |
| | | 4 | 71.94 | 20.14 |
| | | 5 | 87.77 | 50.36 |
| | | 6 | 79.86 | 25.90 |
| | | 7 | 65.47 | 30.94 |
| | | 8 | 76.98 | 1.44 |
| | | 9 | 68.35 | 10.07 |
| | | 10 | 82.01 | 2.16 |
| C_pellans | 7 | 1 | 97.84 | 0.72 |
| | | 2 | 69.78 | 19.42 |
| | | 3 | 99.28 | 0.72 |
| | | 4 | 96.40 | 0.00 |
| | | 5 | 64.03 | 26.62 |
| | | 6 | 45.32 | 20.86 |
| | | 7 | 69.78 | 2.16 |
| C_persimilis | 6 | 1 | 98.56 | 0.72 |
| | | 2 | 99.28 | 13.67 |
| | | 3 | 66.91 | 29.50 |
| | | 4 | 94.96 | 0.00 |
| | | 5 | 63.31 | 29.50 |
| | | 6 | 82.01 | 30.22 |
| C_persimplex | 11 | 1 | 98.56 | 1.44 |
| | | 2 | 77.70 | 22.30 |
| | | 3 | 88.49 | 5.76 |
| | | 4 | 68.35 | 0.00 |
| | | 5 | 98.56 | 1.44 |
| | | 6 | 93.53 | 0.00 |
| | | 7 | 100.00 | 18.71 |
| | | 8 | 84.89 | 6.47 |
| | | 9 | 77.70 | 36.69 |
| | | 10 | 51.08 | 15.83 |
| | | 11 | 90.65 | 21.58 |
| C_pusillus | 5 | 1 | 52.52 | 1.44 |
| | | 2 | 99.28 | 5.76 |
| | | 3 | 95.68 | 1.44 |
| | | 4 | 79.86 | 26.62 |
| | | 5 | 63.31 | 21.58 |
| C_rohweri | 22 | 1 | 98.56 | 0.00 |
| | | 2 | 77.70 | 0.72 |
| | | 3 | 93.53 | 0.00 |
| | | 4 | 76.26 | 19.42 |
| | | 5 | 99.28 | 28.78 |

| | | | | |
|---|---|---|---|---|
| | | 6 | 79.86 | 2.16 |
| | | 7 | 85.61 | 1.44 |
| | | 8 | 85.61 | 69.06 |
| | | 9 | 76.98 | 47.48 |
| | | 10 | 96.40 | 0.00 |
| | | 11 | 99.28 | 1.44 |
| | | 12 | 82.73 | 25.18 |
| | | 13 | 97.84 | 0.72 |
| | | 14 | 66.91 | 31.65 |
| | | 15 | 61.15 | 18.71 |
| | | 16 | 82.73 | 0.72 |
| | | 17 | 64.75 | 28.78 |
| | | 18 | 50.36 | 27.34 |
| | | 19 | 46.76 | 17.27 |
| | | 20 | 35.19 | 38.27 |
| | | 21 | 39.57 | 9.35 |
| | | 22 | 48.92 | 16.55 |
| C_similimus | 6 | 1 | 94.24 | 3.60 |
| | | 2 | 97.12 | 14.39 |
| | | 3 | 64.75 | 0.00 |
| | | 4 | 85.61 | 2.88 |
| | | 5 | 64.03 | 20.86 |
| | | 6 | 80.58 | 33.09 |
| C_spinosus | 8 | 1 | 94.96 | 2.88 |
| | | 2 | 87.77 | 8.63 |
| | | 3 | 72.66 | 12.95 |
| | | 4 | 41.73 | 3.60 |
| | | 5 | 98.56 | 0.00 |
| | | 6 | 58.99 | 22.30 |
| | | 7 | 99.28 | 3.60 |
| | | 8 | 89.93 | 1.44 |
| C_umbraculatus | 14 | 1 | 2.47 | 0.00 |
| | | 2 | 74.10 | 28.06 |
| | | 3 | 92.81 | 10.79 |
| | | 4 | 99.28 | 1.44 |
| | | 5 | 99.28 | 27.34 |
| | | 6 | 96.40 | 13.67 |
| | | 7 | 82.01 | 11.51 |
| | | 8 | 85.61 | 18.71 |
| | | 9 | 94.96 | 6.47 |
| | | 10 | 97.12 | 16.55 |
| | | 11 | 69.78 | 5.04 |
| | | 12 | 74.10 | 30.22 |
| | | 13 | 98.56 | 1.44 |
| | | 14 | 41.01 | 6.47 |
| _varians_PL005 | 11 | 1 | 97.84 | 3.60 |
| | | 2 | 93.53 | 37.41 |
| | | 3 | 77.70 | 42.45 |
| | | 4 | 99.28 | 0.72 |
| | | 5 | 93.53 | 0.00 |
| | | 6 | 75.54 | 32.37 |
| | | 7 | 58.27 | 39.57 |

| | | | | |
|---|---|---|---|---|
| | | 8 | 98.56 | 24.46 |
| | | 9 | 82.01 | 61.15 |
| | | 10 | 87.05 | 29.50 |
| | | 11 | 41.73 | 23.02 |
| _varians_PL010 | 11 | 1 | 87.79 | 4.82 |
| | | 2 | 89.69 | 3.42 |
| | | 3 | 95.41 | 3.46 |
| | | 4 | 97.39 | 3.64 |
| | | 5 | 96.40 | 2.72 |
| | | 6 | 96.76 | 5.83 |
| | | 7 | 98.87 | 4.82 |
| | | 8 | 96.12 | 5.00 |
| | | 9 | 94.58 | 4.38 |
| | | 10 | 96.62 | 1.81 |
| | | 11 | 90.69 | 2.10 |

| Number contigs bin | Number genes bin | Length bin (Mb) | Number of BGC present |
|---|---|---|---|
| 316 | 1,506 | 1.23 | 0 |
| 66 | 2,075 | 2.55 | 1 |
| 3,112 | 8,646 | 11.92 | 0 |
| 2,760 | 8,783 | 10.34 | 0 |
| 323 | 2,509 | 2.22 | 0 |
| 38 | 1,402 | 1.52 | 0 |
| 176 | 2,053 | 2.21 | 0 |
| 729 | 5,094 | 4.73 | 0 |
| 275 | 2,115 | 1.89 | 0 |
| 85 | 2,053 | 2.14 | 0 |
| 455 | 3,733 | 3.50 | 1 |
| 910 | 4,862 | 4.23 | 1 |
| 798 | 4,765 | 4.39 | 1 |
| 523 | 2,575 | 2.21 | 1 |
| 136 | 2,006 | 2.42 | 0 |
| 425 | 2,743 | 2.77 | 0 |
| 194 | 423 | 0.542 | 0 |
| 525 | 4,276 | 4.48 | 0 |
| 632 | 5,449 | 5.14 | 1 |
| 717 | 4,724 | 4.39 | 1 |
| 143 | 1,824 | 2.03 | 0 |
| 508 | 2,165 | 2.26 | 0 |
| 805 | 3,940 | 4.53 | 0 |
| 448 | 2,497 | 2.80 | 0 |
| 299 | 2,013 | 1.76 | 0 |
| 181 | 3,023 | 3.06 | 1 |
| 640 | 4,304 | 3.73 | 1 |
| 71 | 2,152 | 2.61 | 0 |
| 588 | 4,057 | 3.90 | 1 |
| 526 | 2,793 | 2.28 | 0 |
| 736 | 4,462 | 3.88 | 2 |
| 79 | 1,817 | 1.99 | 0 |
| 523 | 2,225 | 2.23 | 0 |
| 90 | 2,357 | 2.46 | 0 |
| 871 | 5,889 | 5.54 | 5 |
| 672 | 4,083 | 4.01 | 0 |
| 490 | 3,821 | 3.76 | 0 |
| 525 | 3,808 | 3.56 | 1 |
| 222 | 1,103 | 0.94 | 0 |
| 338 | 1,693 | 1.47 | 0 |
| 398 | 3,704 | 3.99 | 2 |
| 832 | 4,229 | 4.03 | 2 |
| 635 | 4,010 | 3.33 | 2 |
| 472 | 2,911 | 3.57 | 0 |
| 290 | 3,308 | 3.44 | 0 |
| 1,300 | 6,793 | 6.04 | 2 |
| 26 | 2,147 | 2.54 | 1 |
| 766 | 3,583 | 3.12 | 1 |
| 176 | 2,383 | 2.40 | 0 |
| 352 | 2,530 | 2.37 | 0 |

| | | | |
|---|---|---|---|
| 390 | 2,671 | 3.01 | 1 |
| 328 | 1,740 | 1.44 | 1 |
| 344 | 2,329 | 2.10 | 0 |
| 74 | 2,393 | 2.50 | 1 |
| 436 | 2,528 | 2.28 | 1 |
| 116 | 2,170 | 2.51 | 1 |
| 1,072 | 5,110 | 4.38 | 1 |
| 253 | 1,840 | 1.97 | 0 |
| 533 | 2,760 | 2.94 | 1 |
| 162 | 2,736 | 2.79 | 0 |
| 993 | 4,940 | 4.99 | 0 |
| 1,300 | 6,840 | 6.16 | 6 |
| 472 | 3,815 | 3.76 | 0 |
| 451 | 2,463 | 2.12 | 1 |
| 281 | 2,444 | 2.33 | 0 |
| 533 | 2,918 | 2.63 | 0 |
| 322 | 2,522 | 2.84 | 0 |
| 403 | 2,303 | 2.37 | 0 |
| 918 | 2,904 | 3.10 | 1 |
| 131 | 2,172 | 2.60 | 0 |
| 36 | 4,504 | 4.12 | 0 |
| 825 | 2,801 | 2.58 | 2 |
| 845 | 4,280 | 3.92 | 1 |
| 454 | 4,452 | 4.02 | 0 |
| 475 | 5,164 | 5.07 | 0 |
| 623 | 3,314 | 3.54 | 3 |
| 722 | 2,148 | 2.52 | 0 |
| 22 | 4,195 | 3.49 | 1 |
| 1,261 | 3,784 | 3.03 | 2 |
| 810 | 6,564 | 5.66 | 0 |
| 121 | 1,836 | 1.94 | 0 |
| 1,052 | 3,906 | 4.10 | 0 |
| 1,293 | 4,905 | 5.90 | 0 |
| 189 | 1,467 | 1.33 | 0 |
| 192 | 2,020 | 2.11 | 0 |
| 207 | 2,856 | 2.81 | 1 |
| 497 | 4,608 | 4.34 | 2 |
| 437 | 2,707 | 2.75 | 0 |
| 825 | 4,921 | 4.78 | 0 |
| 556 | 2,557 | 2.28 | 0 |
| 605 | 3,858 | 3.56 | 1 |
| 281 | 3,727 | 3.59 | 0 |
| 429 | 2,381 | 2.73 | 1 |
| 165 | 3,194 | 2.82 | 1 |
| 635 | 3,009 | 2.54 | 0 |
| 635 | 1,228 | 1.15 | 1 |
| 29 | 1,583 | 1.64 | 0 |
| 921 | 6,274 | 8.68 | 0 |
| 47 | 1,729 | 1.82 | 0 |
| 3,378 | 18,614 | 31.63 | 1 |
| 147 | 3,052 | 3.22 | 3 |

| | | | |
|---|---|---|---|
| 59 | 2,164 | 2.42 | 0 |
| 108 | 2,148 | 2.24 | 0 |
| 313 | 4,518 | 4.46 | 1 |
| 424 | 5,174 | 5.36 | 0 |
| 77 | 2,295 | 2.36 | 0 |
| 31 | 2,004 | 2.15 | 0 |
| 202 | 3,421 | 3.46 | 2 |
| 88 | 2,033 | 2.11 | 0 |
| 121 | 2,045 | 2.11 | 1 |
| 131 | 2,775 | 2.87 | 0 |
| 90 | 2,073 | 2.16 | 0 |
| 436 | 2,152 | 2.00 | 0 |
| 323 | 1,693 | 1.48 | 0 |
| 671 | 3,489 | 3.16 | 0 |
| 14,539 | 46,227 | 71.17 | 1 |
| 620 | 2,492 | 1.92 | 0 |
| 13,054 | 31,045 | 45.58 | 0 |
| 328 | 2,123 | 1.99 | 0 |
| 946 | 4,107 | 4.89 | 0 |
| 371 | 1,964 | 1.67 | 0 |
| 275 | 2,249 | 2.34 | 0 |
| 803 | 4,703 | 4.10 | 2 |
| 826 | 4,667 | 3.95 | 2 |
| 50 | 2,077 | 2.53 | 0 |
| 450 | 3,359 | 3.05 | 2 |
| 267 | 2,519 | 2.31 | 1 |
| 173 | 2,059 | 1.90 | 1 |
| 81 | 2,658 | 2.72 | 2 |
| 1,067 | 5,996 | 5.08 | 3 |
| 514 | 4,141 | 4.46 | 1 |
| 825 | 3,903 | 4.28 | 1 |
| 1,298 | 2,744 | 3.83 | 0 |
| 664 | 3,383 | 3.26 | 1 |
| 332 | 4,114 | 3.83 | 3 |
| 147 | 2,493 | 2.45 | 1 |
| 4,233 | 11,471 | 18.38 | 0 |
| 3,922 | 11,620 | 16.85 | 0 |
| 2,918 | 8,842 | 11.08 | 0 |
| 669 | 3,307 | 4.09 | 2 |
| 753 | 4,373 | 5.11 | 2 |
| 158 | 2,106 | 2.13 | 0 |
| 112 | 1,455 | 1.26 | 0 |
| 374 | 2,580 | 2.58 | 1 |
| 109 | 1,957 | 2.27 | 0 |
| 5,345 | 12,816 | 17.88 | 0 |
| 1,020 | 4,203 | 4.60 | 0 |
| 211 | 2,449 | 2.52 | 0 |
| 459 | 3,650 | 3.60 | 1 |
| 253 | 2,957 | 2.90 | 2 |
| 67 | 1,793 | 1.84 | 1 |
| 219 | 3,237 | 3.42 | 2 |
| 224 | 2,656 | 2.74 | 0 |

| | | | |
|---|---|---|---|
| 120 | 2,542 | 2.65 | 0 |
| 422 | 3,599 | 3.55 | 1 |
| 583 | 3,581 | 3.55 | 2 |
| 480 | 3,651 | 3.53 | 0 |
| 142 | 1,237 | 1.65 | 0 |
| 136 | 1,842 | 1.70 | 0 |
| 92 | 1,438 | 2.01 | 0 |
| 62 | 1,983 | 2.01 | 0 |
| 45 | 2,475 | 2.04 | 0 |
| 126 | 3,585 | 2.91 | 0 |
| 112 | 1,586 | 2.30 | 0 |
| 101 | 2,542 | 2.16 | 0 |
| 84 | 2,079 | 1.99 | 0 |
| 87 | 1,972 | 1.85 | 0 |
| 63 | 1,845 | 1.49 | 0 |
